# The BR-body proteome contains a complex network of protein-protein and protein-RNA interactions

**DOI:** 10.1101/2023.01.18.524314

**Authors:** Nandana V., Rathnayaka-Mudiyanselage I.W., Muthunayake N.S., Hatami A., Mousseau C.B., Ortiz-Rodríguez L.A., Vaishnav J., Collins M., Gega A., Mallikaarachchi K.S., Yassine H., Ghosh A., Biteen J.S., Zhu Y., Champion M.M., Childers W.S., Schrader J.M.

## Abstract

Bacterial RNP bodies (BR-bodies) are non-membrane-bound structures that facilitate mRNA decay by concentrating mRNA substrates with RNase E and the associated RNA degradosome machinery. However, the full complement of proteins enriched in BR-bodies has not been defined. Here we define the protein components of BR-bodies through enrichment of the bodies followed by mass spectrometry-based proteomic analysis. We found 111 BR-body enriched proteins, including several RNA binding proteins, many of which are also recruited directly to *in vitro* reconstituted RNase E droplets, showing BR-bodies are more complex than previously assumed. While most BR-body enriched proteins that were tested cannot phase separate, we identified five that undergo RNA-dependent phase separation *in vitro*, showing other RNP condensates interface with BR-bodies. RNA degradosome protein clients are recruited more strongly to RNase E droplets than droplets of other RNP condensates, implying that client specificity is largely achieved through direct protein-protein interactions. We observe that some RNP condensates assemble with preferred directionally, suggesting that RNA may be trafficked through RNP condensates in an ordered manner to facilitate mRNA processing/decay, and that some BR-body associated proteins have the capacity to dissolve the condensate. Finally, we find that RNA dramatically stimulates the rate of RNase E phase separation *in vitro*, explaining the dissolution of BR-bodies after cellular mRNA depletion observed previously. Altogether, these results suggest that a complex network of protein-protein and protein-RNA interactions controls BR-body phase separation and RNA processing.

**Highlights:**

1. BR-body proteomics identified 111 proteins enriched in BR-bodies.
2. BR-bodies associate with an interconnected network of RNP condensates.
3. BR-body condensation is modulated by its interaction network.
4. RNA is required for rapid BR-body condensation.

**Graphical Abstract:** Summary of the BR-body protein interactome. Lines between two protein circles represent a direct interaction.

## Introduction

Biomolecular condensates have been shown to be an important mode of subcellular organization in eukaryotes and bacteria^1–3^. As bacteria often lack membrane-bound organelles, biomolecular condensates may provide a generalized strategy for their subcellular organization. Indeed, many bacterial biomolecular condensates have been recently identified in various species, including BR-bodies, signaling condensates, DNA replication condensates, RNApol condensates, Hfq condensates, Rubisco condensates, etc.^4–10^. Biomolecular condensates typically assemble through the physical process of phase separation, which involves association of macromolecular scaffolds and segregation of other cellular components, ultimately leading to distinct non-membrane bound structures^11^. Recent analyses of condensate physical properties appear to be consistent with a model of percolation-coupled phase separation, which includes both associative interactions as well as segregative^12^, leading to phase separation^13^. Macromolecules that have the capacity to phase separate are referred to as scaffolds, which generally provide a platform for the multi-valent interactions that lead to phase separation. Molecules that are recruited to condensates but cannot phase separate themselves are called clients^14^. A condensate’s composition of scaffolds and clients determines its biochemical and cellular function. Importantly, molecules inside of condensates often show diffusive dynamics, suggesting biomolecular condensates facilitate the biochemical processes within them^15, 16^. Several models have been proposed for how condensates might accelerate enzyme-catalyzed processes, including increasing reaction rates by raising the local concentration of substrates and enzymes, increasing reaction specificity by the selective recruitment of certain substrates and avoidance of others, and increasing reaction completion by preventing the release of pathway intermediates^17–19^. Conversely, biomolecular condensates have also been proposed to sequester substrate molecules from their enzymes, leading to a reduction in reaction rates^20, 21^. Therefore, it is important to identify the essential molecular components of the condensate to determine the biochemical consequences of biomolecular condensate organization.

Despite advances in biomolecular condensate biochemistry, many *in vitro* studies have focused on purified scaffolds, and often lack the full suite of molecules identified to interact *in vivo*. This is potentially problematic, as proteomic/transcriptomic investigations in eukaryotes have revealed that biomolecular condensates are enriched in hundreds of different molecular species that may alter their biochemical or cellular functions. For example, P-bodies have been found to contain >125 proteins and >6000 RNA transcripts^22^, and importantly appear to help concentrate mRNA decay enzymes (including decapping enzymes, LSM proteins, Xrn1 nuclease, deadenylases, etc.) together with decapped and translationally repressed mRNAs. Stress granules have been found to contain >300 proteins and >1800 RNA transcripts^23–25^, often including a partially overlapping set of components with P-bodies^22, 26^, yet the differences in composition and slower dynamics^27^ suggests that these structures likely have distinct functions. Through careful quantitative measurements, the core set of P-body proteins has recently been reconstituted *in vitro*, with similar dynamic properties^28^. In bacteria, BR-bodies were found to be enriched in long, poorly translated cellular mRNAs and sRNAs, a class of translational repressors, together with the RNA degradosome proteins^4, 29^. Therefore, it was assumed that this structure might have a small set of protein clients that could be amendable to full *in vitro* reconstitution ^27^; however, the full complement of proteins in BR-bodies has not been identified.

To define the proteome of *Caulobacter crescentus* BR-bodies, we performed cellular enrichment by differential centrifugation^30^ followed by quantitative liquid chromatography tandem mass spectrometry (LC-MS). Similar to eukaryotic condensates, we identified over 100 proteins that were enriched in BR-bodies. Enriched proteins tended to localize into foci *in vivo* and enter RNase E condensates *in vitro*, suggesting many are likely direct clients of RNase E, the main scaffold of the BR-body. While most enriched proteins do not assemble into biomolecular condensates *in vitro*, we identified five BR-body associated proteins that can undergo phase-separation (phase separate without RNase E), suggesting they form distinct RNP condensates in the cell that interact with RNase E. By assaying the pairwise ability of each scaffold protein to mix with the others, we observed that some pairs of scaffolds have strong associations, and that certain pairs show preferred directional recruitment, which likely impacts RNP assembly order, leading to heterogenous RNP condensates *in vivo*. We also identified two proteins that can dissolve BR-bodies both *in vivo* and *in vitro*, suggesting that they may negatively regulate the cell’s ability to assemble BR-bodies. Despite the identification of many BR-body associated proteins, the RNA degradosome proteins are the only proteins identified that stoichiometrically bind to RNase E^31^ and that are uniformly colocalized with RNase E foci *in vivo*, suggesting a minimal BR-body may be possible to reconstitute. Minimal BR-body phase separation was kinetically stimulated by RNA and showed similar RNase E FRAP dynamics to RNase E-RNA droplets. Overall, this suggests that BR-bodies have similar complex interaction network to eukaryotic RNP condensates, and the reconstitution of the “core” minimal BR-body components provides an exciting framework to begin assessing their biochemical functions and how they might be altered by the addition of BR-body associated proteins.

## Results

### Characterization of the BR-body proteome by LC-MS Proteomics

To define the *C. crescentus* BR-body associated proteome, BR-body enrichment was performed as in^30^ and subjected to bottom-up proteomics preparation and analysis (Fig 1A, Table S1). BR-body enriched samples were compared to a mock treated lysate of a mutant that is unable to assemble BR-bodies due to the deletion of the C-terminal IDR of RNase E^30^. 100 μg protein was prepared in triplicate for either condition and converted into peptides via enzymatic digestion using S-Traps^32^. Each sample was analyzed by LC-MS in technical triplicate. Peptide-spectral matching, protein inference, and Label Free Quantification (LFQ) were performed using FlashLFQ in MetaMorpheus^33^. Data were filtered to an FDR of 1.0% and exported as text for analysis. Using these parameters, a total of 1079 proteins were identified in both samples, and the ratio of signal in the BR-body enriched sample (JS299) over the negative control (JS221) was plotted (Fig 1B). 111 proteins were identified to be enriched in BR-bodies. These enriched proteins had Log2 BR-body enrichment value of >1 (>2-fold enrichment) with a >95% confidence interval (resulting p-values subject to Benjamini-Hochberg correction) or were identified in all three biological replicates in BR-body enriched samples but were not detected in the negative control. These enriched proteins were predominantly cytoplasmic, while proteins derived from other cellular compartments were depleted, suggesting reasonable enrichment of BR-bodies occurred in the sample. In addition, we observed significantly higher scores for known RNase E associated proteins, while proteins that interact with polar biomolecular condensates were highly depleted (Fig 1B), suggesting that the enrichment was likely not contaminated with all cellular condensates. GO-term analysis of BR-body enriched proteins revealed RNA binding, ATP binding, GTPase activity, and RNA degradation functionalities (Fig 1B).

**Figure 1.**
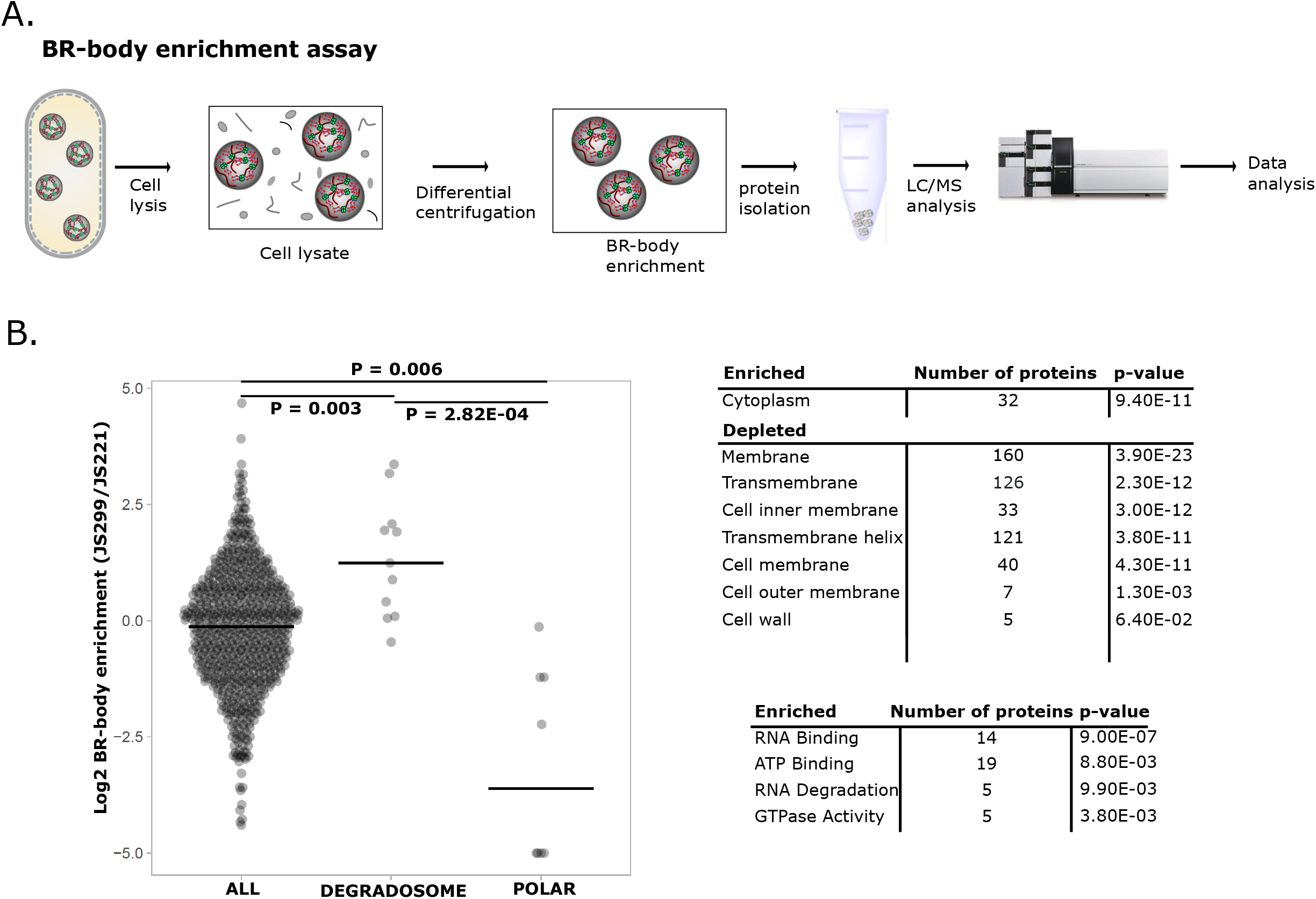
BR-body enrichment proteomics identifies BR-body associated proteins. A) *Caulobacter crescentus* BR-body enrichment^30^ was performed, followed by LC-MS/MS proteomics to identify BR-body associated proteins. JS299 (an RNase E active site mutant) was used to isolate BR-bodies, and JS221 (an RNase E mutant lacking the C-terminal disordered region that cannot form BR-bodies) was used as a negative control. B) BR-body enrichment analysis from the LC-MS/MS proteomics data. The RNA degradosome proteins were identified by pulldown from^31, 34, 35^. Polar proteins were identified as interactors with PopZ^38^ or polar condensates with interactions with PopZ, PodJ, or SpmX. Polar proteins measured in the negative control but undetected in the BR-body enriched samples were plotted at <-5. T-tests with uneven variance were used for statistical comparison. GO-term enrichment for localization and protein function were performed using NCBI DAVID^71^.

To validate whether BR-body enriched proteins localize into BR-bodies *in vivo*, we examined the ability of enriched proteins to localize into foci *in vivo*, the foci’s dependence on RNase E expression, and the proteins’ colocalization with BR-bodies. To comprise the set of BR-body enriched proteins, we chose proteins that are part of the *C. crescentus* RNA degradosome ^31, 34^, proteins that associate with RNase E in the cold ^35^, and proteins that are known to have a role in RNA processes. We selected proteins with no known role in RNA processes (DnaK and FabG) as a negative control. In addition, we also chose a set of proteins that were not enriched in BR-bodies, but function in RNA decay. We expressed each protein with a YFP tag from the vanillate locus (Fig S1) and determined the average number of foci per cell (Fig 2A). Importantly, even when expressed at very high levels in *Caulobacter*, YFP is not sufficient to form foci *in vivo*^36^. Interestingly, 7/12 proteins from the BR-body enriched set assembled into foci in the cells, while only 2/10 proteins (RNase HI and MetK) from the negative control set showed foci. Of the 5/12 proteins that were enriched but did not form foci, it is possible that the C-terminal YFP tag interfered with their ability to form foci, as SmpB, which was found to form foci by immunofluorescence^37^, did not form foci when fused to YFP.

**Figure 2.**
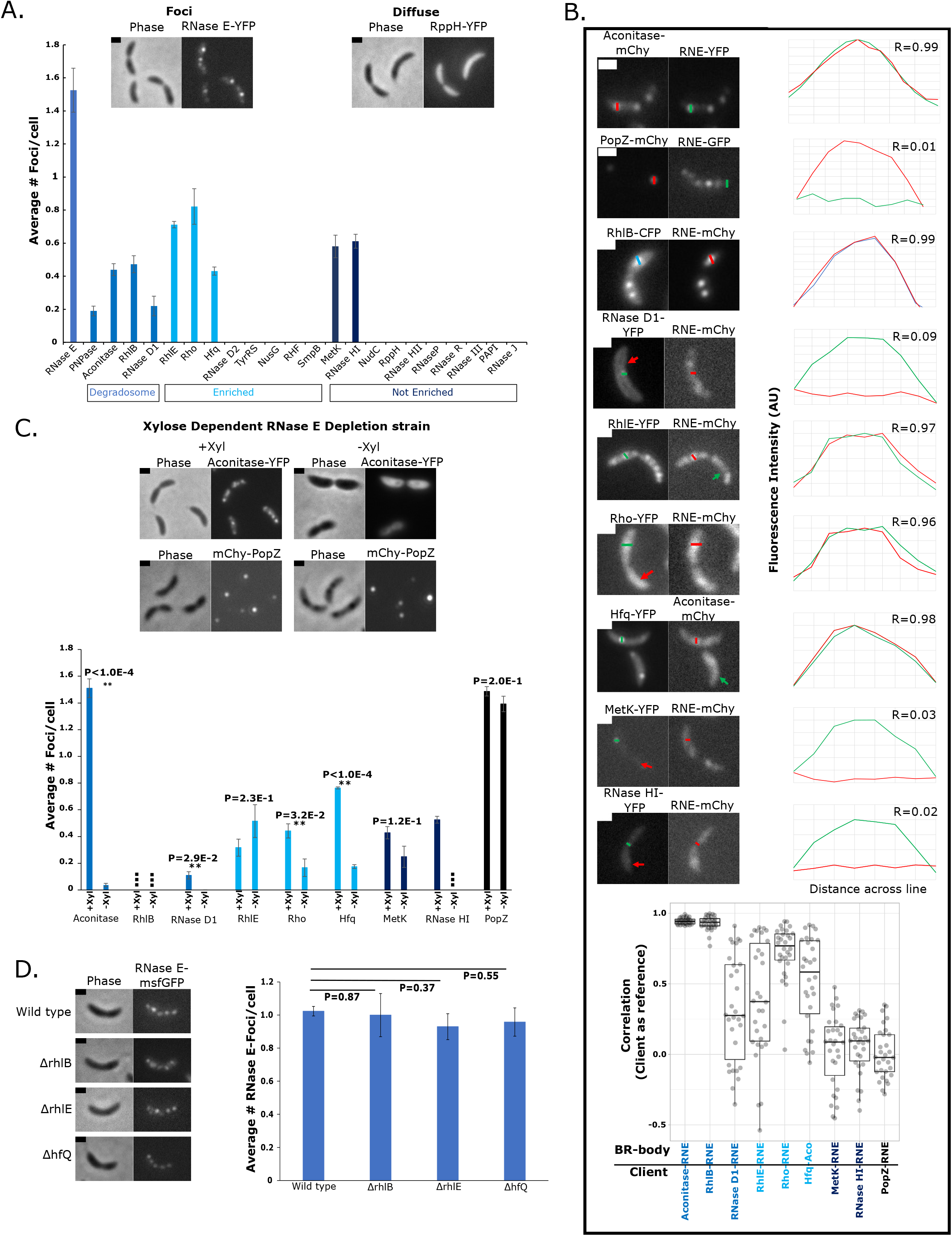
RNase E is required for the foci formation and BR-body colocalization of most BR-body associated proteins. A) The localization pattern of foci-forming YFP fusions. All genes were placed under control of the vanillate promoter and grown in M2G minimal media. *In vivo* protein fusion strains were analyzed at mid-exponential phase of growth (OD 0.3-0.6) following 6-hours of induction with 0.5mM vanillate. and quantified using MicrobeJ^68^. Three replicates were performed on three different days and three images were analyzed from each day to calculate the average number of foci per cell. A minimum of 150 cells were used for each YFP-fusion for the analysis. Error bars represent the standard error. Representative images are in supplementary figure 1 (Fig S1). B) Colocalization patterns of the BR-body enriched protein fusions with BR-body markers. Colored line traces indicate the pattern of localization observed for individual foci. The line trace method was used to assess enrichment. The correlation was calculated for each line trace using the BR-body enriched protein (client) as a reference, and each line trace correlation value was plotted as a single dot on the plot. A minimum of 30 foci for each BR-body enriched protein were used for the analysis. The median value is represented as a black line. Statistics were based upon a two-tailed t-test with unequal variance. C) Subcellular localization patterns of *C. crescentus* protein-YFP strains expressed from the *vanA* locus in the RNase E depletion background where the sole copy of the RNase E gene is controlled by the xylose promoter. The YFP intensity of Aconitase-YFP was too low to measure when expressed from the vanillate promoter, so a native gene fusion was used instead. The YFP intensity of each image was normalized from its brightest level relative to its background level. Depletion strains were analyzed at mid-exponential phase of growth (OD 0.3-0.6) and after 24 hours of xylose depletion. Quantitation of the number of *Ccr* protein-YFP foci per cell measured in M2G minimal media with xylose (+Xyl) and M2G minimal media lacking xylose (-Xyl) for the same strains in Fig S2. The squares represent the strains whose levels were too low to detect. All images were quantified using MicrobeJ^68^. Three replicates were performed on three different days and three images were analyzed from each day to calculate the average number of foci per cell. A minimum of 175 cells were used for the analysis. Error bars represent the standard error. Representative images in supplementary figure 2(Fig S2). Foci/cell calculations were performed as in A. D) RNase E GFP foci per cell were measured in wild type, rhlB, rhlE, and hfq mutant backgrounds. Foci/cell calculations were performed as in A. Error bars represent the standard error from a minimum of 100 cells. Scale bar is 1 µm for all images.

To test whether the observed foci were BR-bodies, we examined the colocalization of foci with either the core BR-body scaffold RNase E or its direct binding partner Aconitase^4^. To examine colocalization, we drew a line across the foci of each protein, and then correlated the intensity between this channel and that of the BR-body marker (Fig 2B). As a negative control, we included PopZ, a protein that can form a polar biomolecular condensate involved in cell signaling ^38^. We could not visualize PNPase-YFP together with RNase E-mCherry or Aconitase-mCherry and we noticed very slow growth rates of this strain. We found that for the core degradosome proteins Aconitase and RhlB ^31^, foci were uniformly colocalized with RNase E in all cases (average Correlation = 0.94 Aconitase, 0.92 RhlB), while RNase E foci did not correlate with PopZ foci (average Correlation = 4.8 × 10^-3^). We found that RNase D1, RhlE, Rho, and Hfq showed significant but heterogenous colocalization with BR-bodies (average Correlation = 0.32 RNase D1, 0.39 RhlE, 0.72 Rho, 0.52 Hfq), while BR-body depleted proteins RNase H1 and MetK showed no correlation (average Correlation = 3.1 × 10^-2^ MetK, 6.1 × 10^-2^ RNase H1), similar to the PopZ control, suggesting they are indeed not enriched in BR-bodies. Interestingly, RNase D1 was found to be RNase E associated^34^ but was depleted in the BR-body proteomics data, while RNase D2 was found to be highly enriched in BR-bodies, suggesting that they may be interchangeable. We observed that RNase D1-YFP appeared to be localized to the inner membrane, suggesting that perhaps it was lost upon the initial membrane pelleting step of the BR-body enrichment procedure. In line with this hypothesis, we observed significant colocalization between RNase D1-YFP and RNase E-mCherry in live cells (Fig 2B). We did not observe foci of RNase D2-YFP (Figs 2A, S1), suggesting that either it is not-enriched in BR-bodies, or that the YFP tag is disrupting its localization. The heterogeneity of RNase D1, RhlE, and Hfq suggests that they are present in a subset of BR-bodies, suggesting that some BR-bodies have different compositions or protein clients.

### Identification of BR-body associated scaffolds that condense independent of RNase E

Since RNase E is necessary and sufficient to form BR-bodies^4^, we tested whether RNase E was required for BR-body enriched proteins to form foci (Fig 2C). Because RNase E is essential, we used an RNase E depletion strain where the sole copy of RNase E is under control of the xylose promoter^4^. While most proteins maintained similar YFP fluorescence in the RNase E depletion strain, RhlB-YFP, Aconitase-YFP, PNPase-YFP, and RNase H1-YFP were expressed at levels that were too low to detect (Figs 2C, S2). To overcome this low Aconitase-YFP expression, we utilized a strain in which Aconitase-YFP is present at its native loci in the same xylose-dependent RNase E depletion strain^4^. While most of the strains would saturate cultures overnight, PNPase-YFP expression was toxic in the depletion strain background, leading to extremely slow cell growth in liquid culture, so we were unable to assay this strain. As a control we included PopZ, which is known to form an independent biomolecular condensate in *C. crescentus*^6^. For Aconitase, RNase D1, Hfq, and Rho, fluorescent foci levels dropped significantly upon RNase E depletion, suggesting that these proteins require RNase E to assemble into foci. Interestingly, for MetK and RhlE, foci levels were unchanged or higher upon RNase E depletion, similar to the PopZ control, suggesting that these two proteins assemble into foci independent of RNase E. To examine whether these enriched proteins impact RNase E phase separation, we measured RNase E-GFP foci in strains where the enriched proteins were deleted. Due to Aconitase, RNase D1, and Rho being essential, we only inserted RNase E-GFP into RhlB and hfq deletion strains, or an RhlE disruption mutant strain. We observed no significant difference in the number of RNase E-GFP foci in these strains (Fig 2D), suggesting that RhlB, RhlE, and Hfq do not impact BR-body phase separation *in vivo*.

As MetK and RhlE proteins were found to form RNase E-independent foci *in vivo*, we tested whether these proteins could directly self-assemble into condensates *in vitro*. In addition to MetK and RhlE, we also included several proteins from the BR-body enrichment dataset and also proteins known to interact with RNase E (Figs 3, S3A). Purified proteins were incubated at their estimated *in vivo* concentrations^39^ with or without 20 ng/µL of *C. crescentus* total RNA. We found that RhlE, DEAD, Hfq, MetK, and FabG all assembled RNA-stimulated condensates *in vitro* (Figs 3A, B), suggesting that these proteins are scaffolds that can drive RNA-dependent condensation. Conversely, most of the proteins tested, including PNPase, Aconitase, RhlB, RNase D1, Ribosomal protein S1, NudC, RppH, Rho, NusG, DnaK, and Tyrosyl tRNA synthetase (TyrRS) did not undergo phase separation regardless of the presence of RNA (Fig S3B). Consistent with the phase-separation of RhlE, DEAD, and Hfq, the homologous proteins from *E. coli* were recently found to assemble into condensates *in vitro* or when overexpressed *in vivo*^7, 40^. To test whether these structures are condensates, we performed high salt and RNase treatment of pre-formed condensates (Fig 3C), and in each case the condensates were strongly dissociated, suggesting multivalent electrostatic interactions with RNA are likely important for driving phase separation, like in RNase E condensates^4^. RhlE, DEAD, and Hfq contain intrinsically disordered regions (IDRs) (Fig 3A) that have been characterized as important for their *E. coli* homologues’ ability to form droplets^40, 41^. Prior experiments also showed that the IDR of RNase E was both necessary and sufficient for phase separation^4^. To test the function of these IDRs on phase-separation, we deleted the IDRs of *C. crescentus* RhlE, DEAD, and Hfq and tested their ability to phase-separate. We found that for Hfq and DEAD, the IDR deletions showed reduced phase separation, while for RhlE the IDR deletion was indistinguishable from wild-type (Fig S3C). We did not include FabG and MetK in these experiments as no disordered regions were predicted in their sequences (Fig 3A). Overall, this suggests that for some scaffold proteins, IDRs positively influence but are not strictly required for phase separation.

**Figure 3.**
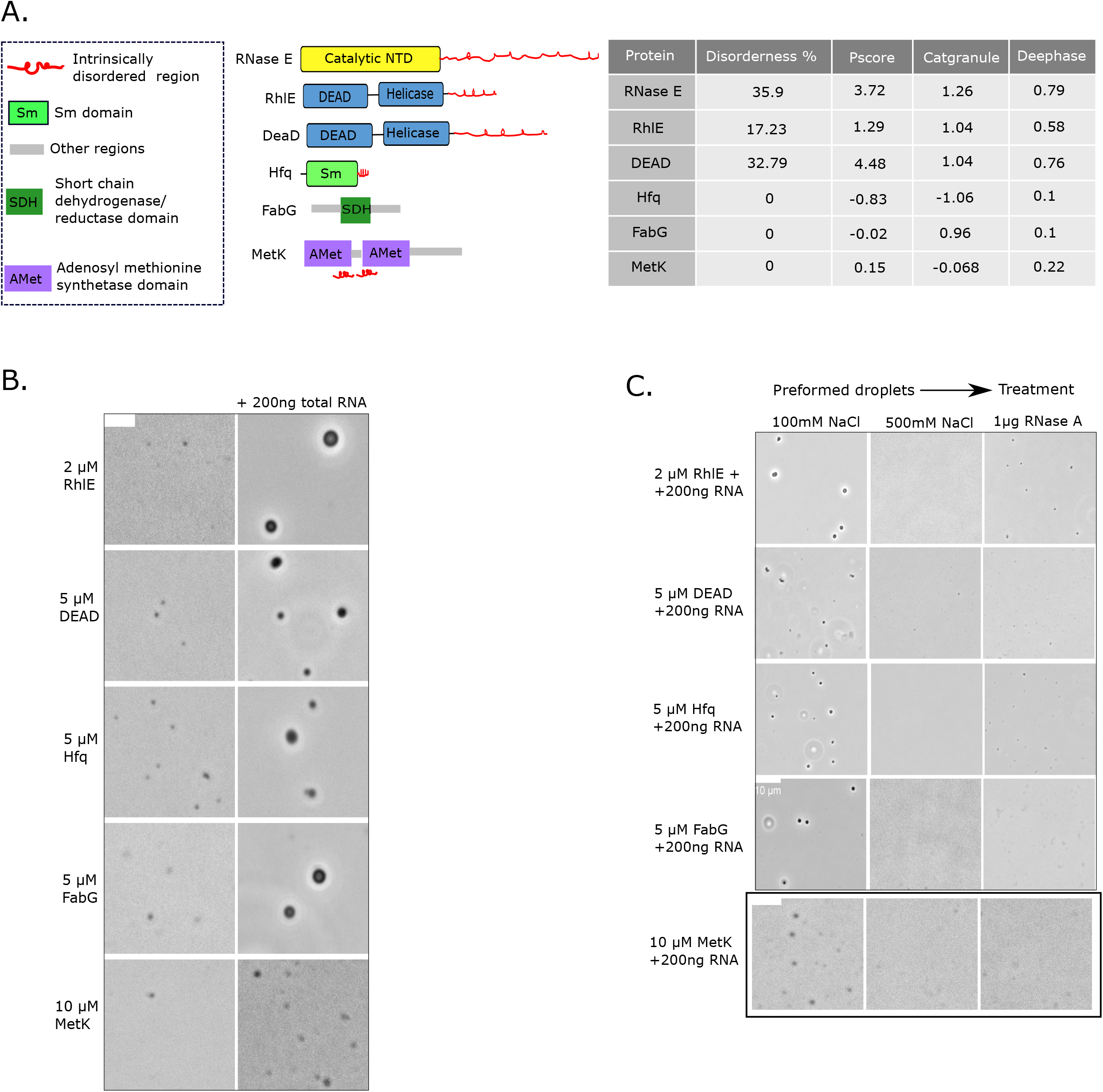
Some BR-body enriched proteins can drive RNA-dependent biomolecular condensation. A) (Left) Domain organization of proteins that undergo LLPS. The red line indicates disordered regions as predicted by PONDR. (Right) Prediction of the propensity to LLPS by Disorderness^72^, Pscore^67^, Catgranule^73^ and DeePhase^62^ scores. B) Phase contrast microscopy images of the purified proteins incubated at their *in vivo* concentrations (DEAD (5 µM), Hfq (5 µM), RhlE (2 µM), FabG (5 µM), MetK (10 µM)) in standard buffer (20mM Tris pH 7.4, 100mM NaCl, 1mM DTT) in the presence and absence of *C. crescentus* total RNA (200 ng). Scale bar is 3µm for all images. C) Droplets are dissolved by NaCl and RNase A. The dissolution of droplets was tested by adding either 0.5M NaCl or RNase A (1 µg) to the preformed droplets. All of the proteins were found to be NaCl and RNase sensitive.

### A complex network of molecular interactions regulates RNP condensation

To better understand which proteins can enter BR-bodies, we selected a set of 18 proteins (Fig S3A) to test further for their recruitment directly into RNase E droplets (Fig 4A). We picked proteins enriched from the BR-body proteomics dataset with known RNA processing roles, proteins known to have protein-protein interactions with RNase E but not detected in the BR-body proteomics experiment, as well as negative control proteins with low BR-body enrichment and no known/predicted interaction with RNase E. In these experiments RNase E and RNA were mixed to form an RNase E condensate, followed by incubation with a Cy5-labeled test protein. The partition coefficient was calculated based on the Cy5 fluorescence (Figs 4B, S4A). We observed that the core RNA degradosome proteins PNPase, Aconitase, RhlB, and RNase D1 all had positive partition coefficients ranging from 2.7 to 4.7, showing that they enter RNase E condensates, while our negative controls BSA and SpmX, which forms a polar signaling condensate^6^, did not enter the RNase E condensate and yielded a partition coefficient near 1, indicating no enrichment. We also identified partition coefficients at similar levels to the core degradosome proteins for RhlE, DnaK, and FabG. In contrast, NudC, RppH, Rho, NusG, and TyrRS all had partition coefficients close to 1, similar to the negative controls BSA and SpmX (Fig 4B). The low partition coefficients for RppH and NudC are consistent with their lack of enrichment in the BR-body proteomics assay; however, Rho, NusG, and TyrRS all had significant enrichment in the BR-body proteomics data, suggesting that they may enter BR-bodies indirectly through another protein. Despite the finding that many proteins were able to enter BR-bodies, none of these proteins altered the partition coefficient of RNase E (Fig S4B), suggesting that most BR-body associated proteins tested did not dramatically influence RNase E’s condensation.

**Figure 4.**
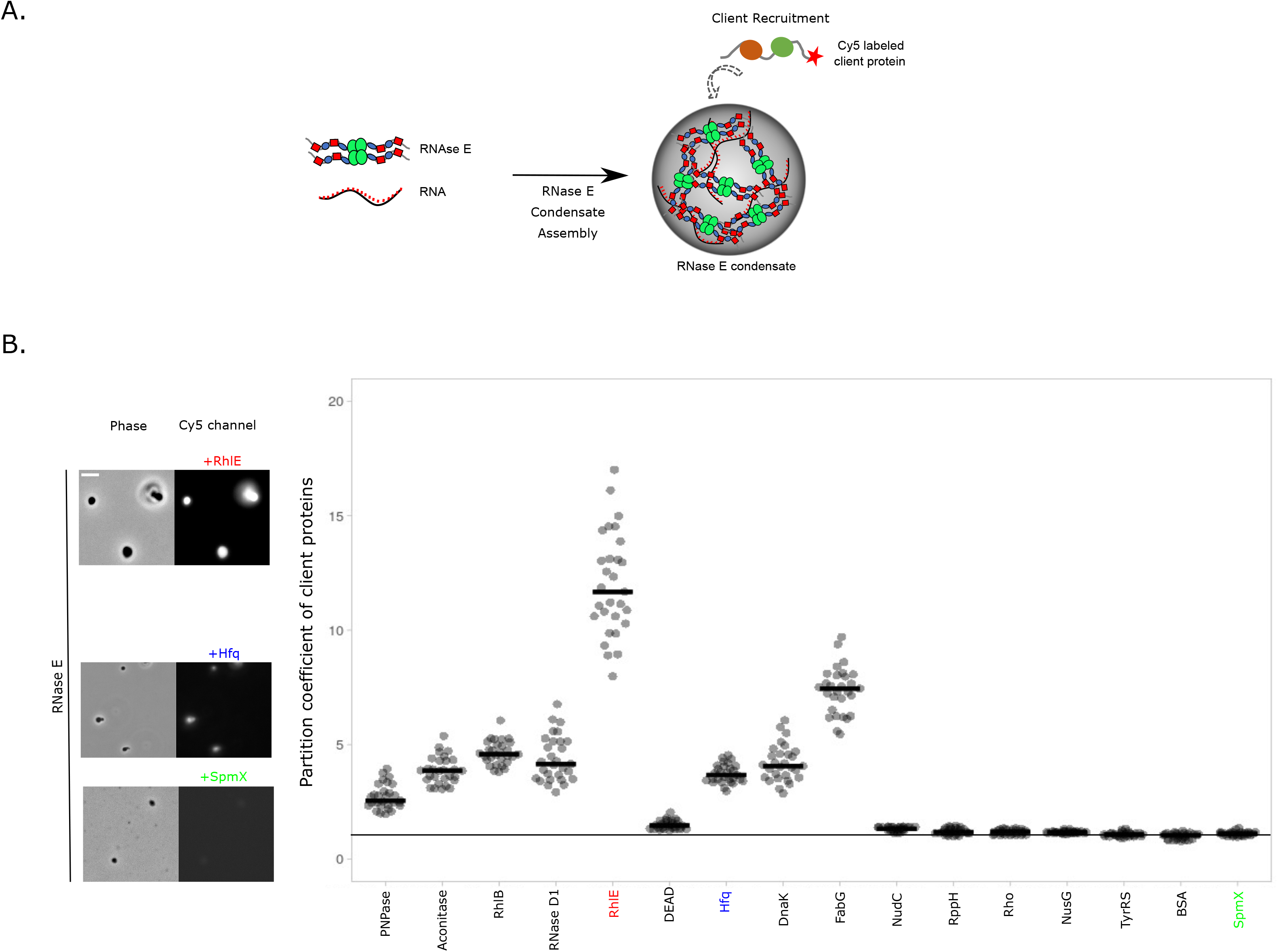
Many BR-body associated proteins are directly recruited to RNase E condensates *in vitro.* A) Cartoon showing the *in vitro* RNase E droplet recruitment assay. Cy5 labeled proteins are added to preformed RNase E CTD+RNA droplets. Microscopy images of the droplets are taken in phase contrast and Cy5 fluorescence, and the partition coefficient of the fluorescence signal is calculated for each droplet. B) Partition coefficient (PC) of Cy5 labeled client proteins in RNase E CTD (6 µM) + total RNA (200ng) droplets. Negative controls BSA-Cy5 and SpmX-Cy5 have partition coefficients of 1. The black horizontal line through the plot indicates a PC of 1, which is the “no enrichment” baseline. PC of ribosomal protein S1 and MetK proteins could not be determined because of the dissolution of droplets. The data were quantified in ImageJ software and plotted in PlotsofData^69^. For the example images presented, the left image is phase contrast and the right image is the Cy5 channel of each indicated protein. The scale is 10 µm for all images. Concentrations of proteins used to quantify the partition coefficient: PNPase (5 µM), Aconitase (5 µM), RhlB (4 µM), RND (5 µM), DEAD(5 µM), Hfq (5 µM), DnaK (5 µM), FabG (5 µM), NudC (2.5 µM), RppH (1 µM), Rho (5 µM), NusG (5 µM), TyrRS (5 µM), and SpmX (5 µM). BSA was added at a 5 µM concentration.

Next, we tested which BR-body associated proteins might induce condensation in the absence of RNA (Fig S5A). We assessed the ability of BR-body associated proteins to stimulate RNase E condensation at their *in vivo* concentrations^39^. PNPase, RhlB, and RhlE stimulated RNase E condensates rather robustly in the absence of RNA (Fig S5A). In addition, FabG and Hfq stimulated irregularly shaped condensates in the absence of RNA, while Aconitase, RNase D1, NudC, RppH, Rho, NusG, DnaK, TyrRS, ribosomal protein S1, and MetK did not stimulate RNase E condensation (Fig S5A). This suggests that a subset of BR-body associated proteins can stimulate RNase E condensation independent of RNA.

Interestingly, we observed that the addition of ribosomal protein S1 (RpsA) or MetK, which were not BR-body enriched (Table S1) but were previously found to associate with RNase E via co-immunoprecipitation (co-IP)^35^, could dissolve RNase E droplets *in vitro* (Figs 4A,5A). Ribosomal protein S1 could only dissociate the droplets if the condensates contained total RNA, while MetK showed strong dissolution if RNase E was condensed with RNA or with the protein PNPase. To test whether the differences between S1 and MetK were related to their inherent RNA binding interactions, we examined their apparent binding on 9S rRNA, a known RNase E substrate^31^. Based on native gel shifts, we found that ribosomal protein S1 bound RNA with tighter apparent affinity than MetK (Fig 5B). The tighter apparent affinity with RNA may explain why ribosomal protein S1 preferentially dissolves RNase E condensates in the presence of RNA. We also used an affinity pull-down assay to test MetK and ribosomal protein S1’s association with MBP-RNase E. Indeed, as compared to the core BR-body protein Aconitase, which is thought to stoichiometrically bind to RNase E^30^, we found that MetK showed less pull-down than Aconitase, while ribosomal protein S1 showed no detectable retention (Figs 5C, S5B). Overall, these data suggest that MetK dissociates BR-bodies through interactions with the RNase E protein (Fig 5). Because MetK showed a weak pull-down signal, we tested the pairwise interaction between MetK and RNase E in a bacterial two-hybrid assay where each protein was expressed recombinantly in *E. coli*^42^ and found that MetK and RNase E do interact *in vivo* (Fig 5D). In the bacterial two-hybrid, we found that Aconitase-T18/RNase E-T25 and MetK-T18/RNAse E-T25 interacts similarly to the T18-Zip/T25-Zip positive control, and RNase E-T25/S1-T18 showed no *lacZ* signal, similar to the T18/T25 negative control. To examine whether the observed *in vitro* dissolution effect of S1 and MetK on BR-bodies is relevant *in vivo*, we inserted the ribosomal protein S1 *rpsA* and *metK* genes into the pBXMCS-2 overexpression vector^43^ and transformed the plasmid into a strain harboring a tagged BR-body marker (Aconitase-mCherry)^4^. We observed *in vivo* that both MetK and ribosomal protein S1 overexpression strongly reduced Aconitase-mCherry foci (Fig 5E), confirming that their dissolution of BR-bodies observed *in vitro* can also occur *in vivo*.

**Figure 5.**
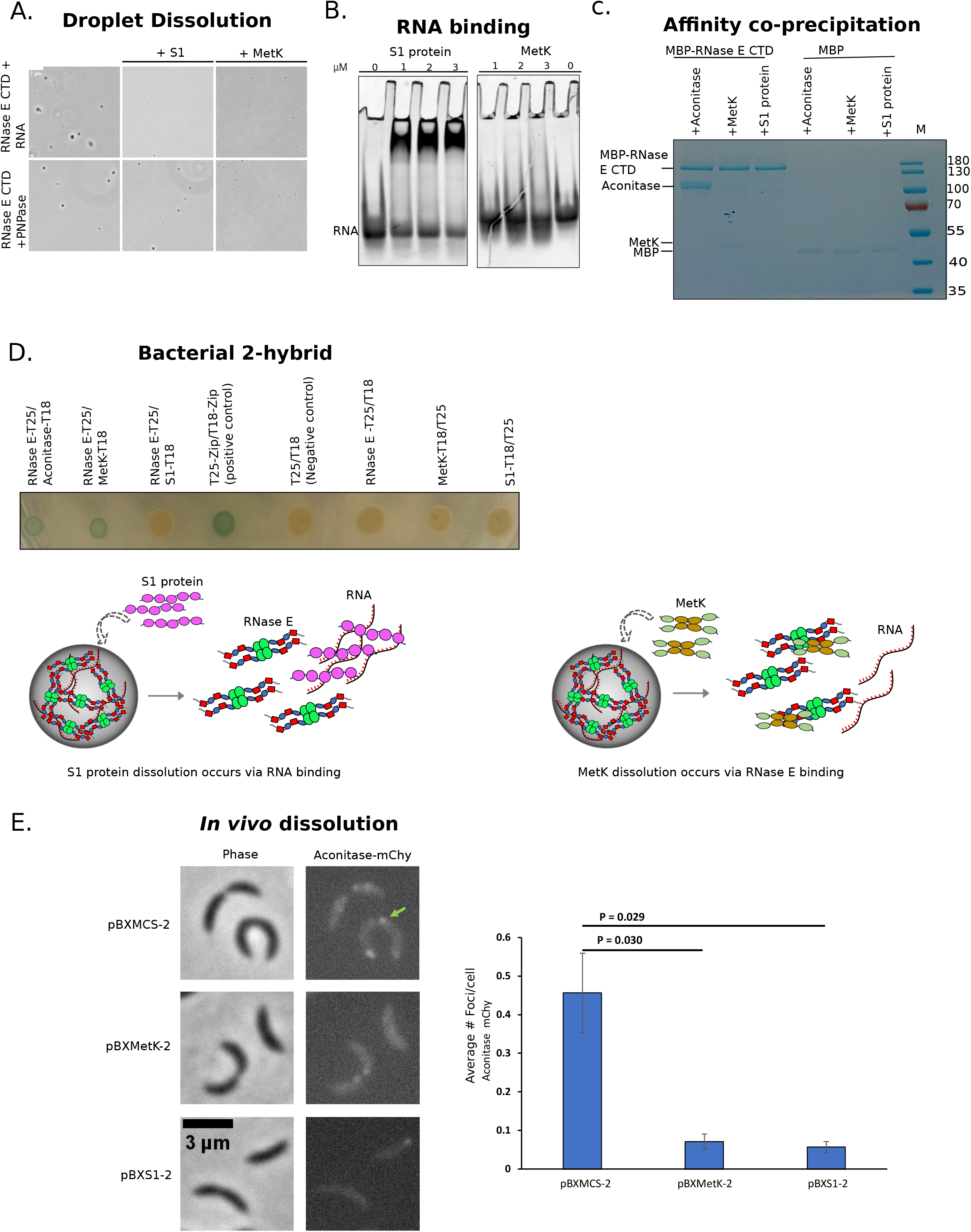
Ribosomal protein S1 and MetK dissolve BR-bodies *in vitro* and *in vivo.* A) Microscopy images of *in vitro* RNase E droplets incubated with either ribosomal protein S1 or MetK. RNA-induced RNase E droplets are shown at the top and PNPase-induced RNase E droplets are shown below. The scale is 10 µm for all images. Concentrations used were: RNase E CTD (10 µM), S1 (5 µM), MetK (5 µM), PNPase (5 µM), and total RNA (20ng/µL). B) RNA and protein binding data for ribosomal protein S1 and MetK with 1 µM RNA. RNA was visualized by SYBR Gold staining. C) Aconitase and MetK pull-down with RNase E. 5 µM RNase E CTD MBP or MBP were incubated with 5 µM of Aconitase or 15 µM of MetK or 15 µM of S1 protein. Eluates were resolved on 10% SDS PAGE. Aconitase and MetK pulled down with RNase E CTD MBP, while S1 protein did not. None of the proteins pulled down with MBP alone. D) Bacterial 2 hybrid assay plate showing the spots of BTH101 cells cotransformed with plasmids expressing the proteins as indicated in the figure. Blue colored spots represent an interaction between the two proteins. The pairs RNase E-T25/Aconitase-T18, RNase E-T25/MetK-T18 and positive control pair T25-Zip/T18-Zip showed interactions and the other pairs showed no interaction. Underneath is a proposed cartoon of the RNA dependence of droplet dissolution by S1 and MetK. E) BR-bodies (marked with Aconitase-mCherry) are dissolved upon overexpression of ribosomal protein S1 or MetK from the pBX multicopy plasmid. Quantitation of the average Aconitase-mCherry foci per cell was performed using MicrobeJ and is plotted on the right (error bars represent the standard error). A minimum of 700 cells were used for each analysis.

Some BR-body enriched proteins were able to assemble into RNP condensates independent of RNase E (Fig 3B), but the client specificity of these RNP condensates is unknown. Because the RNA degradosome client proteins are known to be stoichiometrically bound to RNase E via direct protein-protein interaction sites^31, 35^ and show strong colocalization with BR-bodies (Fig 2B), we tested the specificity of each driver/scaffold to each core RNA degradosome protein labeled with Cy5 dye (Fig 6A). We observed a larger partition coefficient for all Cy5 degradosome clients to RNase E condensates as compared to the other scaffolds/drivers (Figs 6A, S5B), suggesting that the degradosome proteins have a specificity for entering RNase E condensates, likely through their direct protein-protein interactions with RNase E.

**Figure 6.**
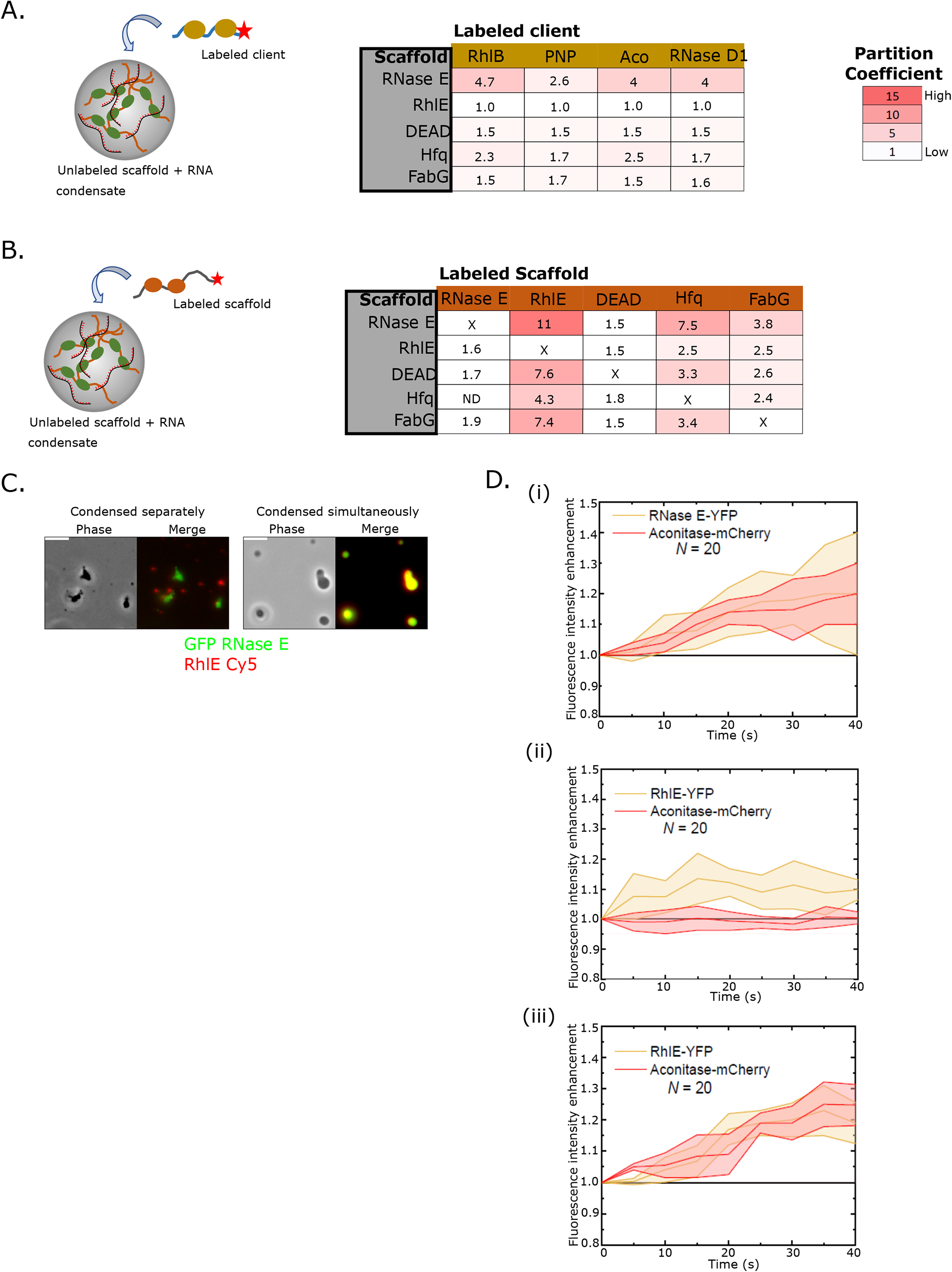
BR-body associated drivers have distinct condensation and client recruitment profiles. A) Scaffold proteins recruit a distinct subset of client proteins. Each RNA degradosome client, labeled with Cy5, was tested for recruitment into pre-formed condensates of each BR-body associated scaffold indicated on the left. BR-body associated scaffolds drive a complex network of dependencies controlling phase separation. Each protein scaffold was incubated with RNA and allowed to phase separate. Then a small concentration of Cy5 labeled proteins were tested for entry into the condensate by measuring the partition coefficients of the Cy5 channel. The partition coefficient was calculated from at least 50 droplets in each case. The data was quantified in ImageJ. Concentrations of the scaffolds used for making condensates were: RNase E (6 µM), RhlE (5 µM), DEAD (5 µM), Hfq (5 µM), and FabG (10µM). Concentrations of the labeled clients added were: RhlB (4 µM), PNP (5 µM), Aconitase (5 µM), and RND (5 µM). B) BR-body associated scaffolds drive a complex network of dependencies controlling phase separation. Each protein scaffold was incubated with RNA and allowed to phase separate. Then a small concentration of Cy5-labeled proteins was tested for entry into the condensate by measuring the partition coefficients of the Cy5 channel. Concentrations of the scaffolds used for making condensates were: RNase E (6 µM), RhlE (5 µM), DEAD (5 µM), Hfq (5 µM), and FabG (10 µM). Concentrations of the labeled scaffolds added were: RNase E (1 µM), RhlE (0.5 µM), DEAD (1 µM), Hfq (1 µM), and FabG (1 µM). Scaffold proteins recruit a distinct subset of client proteins. Each RNA degradosome client labeled with Cy5 was tested for recruitment into each BR-body associated scaffold. Each dot in (A) and (B) plots indicate the partition coefficient in a single droplet. The partition coefficient was calculated from at least 50 droplets in each case. The data were quantified in ImageJ. C) Order of assembly experiments using GFP-RNase E (6 µM) and RhlE-Cy5 (5 µM) condensates. “Condensed simultaneously” indicates that both proteins were added together before the addition of RNA (20ng/µL) to trigger condensation. “Condensed separately” indicates that both proteins were incubated with RNA (20ng/µL) yielding separate condensates that were then mixed together. D) *In vivo* order of foci assembly. Time-lapse two-color imaging of RhlE-YFP and Aconitase-mCherry proteins was performed and the evolution of the intensity of each focus after formation is plotted. (i) Intensity evolution in cells in which both Aconitase-mCherry and RhlE-YFP condensates are formed at approximately at the same time. The middle line indicates the average value and the colored region represents the standard deviation between all replicate events. N is the number of events imaged. (ii) Cells in which RhlE-YFP condensates were formed first and the subsequent appearance of Aconitase-mCherry in RhlE-YFP condensates was tracked. (iii) Cells in which Aconitase-mCherry condensates were formed first and the subsequent appearance of RhlE-YFP in Aconitase-mCherry condensates was tracked.

Next, the miscibility of the drivers/scaffolds of each single RNP condensate was examined (Fig 6B). First, we pre-formed a condensate using an unlabeled scaffold and then added in a Cy5-labeled scaffold in all pairwise combinations and assayed the partition coefficients. We induced condensation in the presence or absence of RNA to test the RNA dependency of each interaction. In the presence of RNA, we assayed condensate formation using the predicted *in vivo* protein concentrations^39^, and in the absence of RNA, condensates were stimulated by increasing the protein concentration above its *in vivo* concentration. For the RNase E condensate, we observed similar recruitment of other scaffolds regardless of whether the condensates were formed with RNA or protein only (Figs 6B, S5C). The partition coefficients for these scaffolds into RNase E condensates correlated with the BR-body enrichment measured previously. RhlE was recruited most strongly into the RNase E condensate (11 partition coefficient) and had the highest observed BR-body enrichment (10 fold). RhlE was followed by Hfq (7.5 partition coefficient, 3.2 BR-body enrichment), FabG (3.8 partition coefficient, 2.2 fold BR-body enrichment), and DEAD (1.5 partition coefficient, 1.6 fold BR-body enrichment). RhlE was recruited strongly into other RNP condensates, with partition coefficients ranging from 4.3 to 11, while the other scaffolds were recruited poorly into RhlE-RNA condensates, with partition coefficients ranging from 1.5 to 2.5. In the absence of RNA, RhlE strongly recruited RNase E (12 partition coefficient) or DEAD (9.8 partition coefficient) while Hfq and FabG maintained similar partition coefficients (Fig S5C). DEAD condensates showed strong recruitment of RhlE and lower recruitment of the other scaffolds, with subtle changes in the presence or absence of RNA. RhlE and FabG were similarly recruited to Hfq condensates regardless of the presence of RNA. RNase E dissolved Hfq-RNA condensates while Hfq protein-only condensates strongly recruited RNase E (6.9 partition coefficient). DEAD was recruited at a low level in Hfq-RNA condensates (1.8 partition coefficient) and dissolved Hfq-only condensates. FabG condensates only formed in the presence of RNA and showed the strongest recruitment of RhlE (7.4 partition coefficient) and Hfq (3.4 partition coefficient). In summary, the scaffold proteins that assemble diverse RNP condensates show specificity of recruitment, suggesting that some RNP condensates can intermix, while others likely do not.

The asymmetric recruitment/dissolution (Fig 6B) and specificity of clients (Fig 6A) observed may help shape the composition of the heterogeneous RNP condensates observed in the cell (Fig 2B). We chose to study the RhlE-RNase E pair further because it showed the strongest directional recruitment in the presence of RNA (Fig 6B). Specifically, RhlE was recruited robustly into RNase E droplets, but RNase E was poorly recruited into RhlE droplets. First, we altered the order of addition of RhlE and RNase E before triggering phase separation (Fig 6C). If the proteins were mixed before condensation with RNA, we observed that RNase E and RhlE mix uniformly. However, when each protein was pre-assembled into a condensate with RNA before mixing both RNP condensates together, we observed that RNase E and RhlE condensates did not readily mix (Fig 6C). This contrasts with the prior observation that free RhlE readily entered RNase E condensates (Fig 6B), suggesting that the order of assembly influences the composition of RNase E and RhlE condensates.

To examine the order of RNase E and RhlE condensate formation *in vivo*, we took time-lapse movies of RhlE and RNase E foci formation in cells. As a positive control, we observed that RNase E-YFP and the core degradosome protein Aconitase-mCherry foci appear simultaneously (Fig 6D, S6A) with uniform colocalization. When BR-bodies appear first, as indicated by Aconitase-mCherry foci, we regularly observed RhlE-YFP signal within these foci (Fig 6D), consistent with the ability of RNase E condensates to recruit RhlE *in vitr*o (Fig 6B). In contrast, RhlE-YFP foci that appeared first generally lacked a colocalized Aconitase-mCherry focus, consistent with the lack of *in vitro* recruitment of RNase E by RhlE (Fig 6B). Taken altogether, these observations suggest that the molecular interactions driving RhlE and RNase E condensation control the ability of these condensates to co-assemble and mix both *in vitro* and *in vivo*. To further test whether the material properties of RhlE and RNase E condensates affect mixing, we performed *in vitro* FRAP on RhlE and RNase E droplets (Fig S6B). We observed that RNase E rapidly recovers either in RNase E-RNA condensates or in mixed RNase E-RhlE-RNA condensates, while RhlE showed little to no recovery in either RhlE-RNA condensates or RNase E-RhlE-RNA condensates (Fig S6B). This suggests that the more dynamic exchange in RNase E droplets compared to RhlE droplets contributes to the preferred directional RhlE to RNase E assembly/mixing observed *in vitro* and *in vivo*.

As RNase E did not appear to readily enter other associated RNP condensates (Fig 6B), we wanted to better understand the role of RNA and the associated degradosome proteins on RNase E condensation. We therefore examined the kinetics of condensation via time-lapse microscopy (Fig 7A, 7B). This assay likely reflects the kinetic processes of phase-separation and surface wetting of the microscope slide. When incubated alone at its predicted *in vivo* concentration, RNase E did not phase separate *in vitro* even after long incubations of many minutes. The addition of RNA strongly stimulated the rate of condensation, occurring on the sub-minutes timescale, consistent with the known importance of RNA for RNase E condensation *in vivo* and in end-point *in vitro* assays^4, 29^. When RNase E was incubated at concentrations much higher than its *in vivo* concentration, it was able to phase separate without RNA after many minutes, (Fig S7).

**Figure 7.**
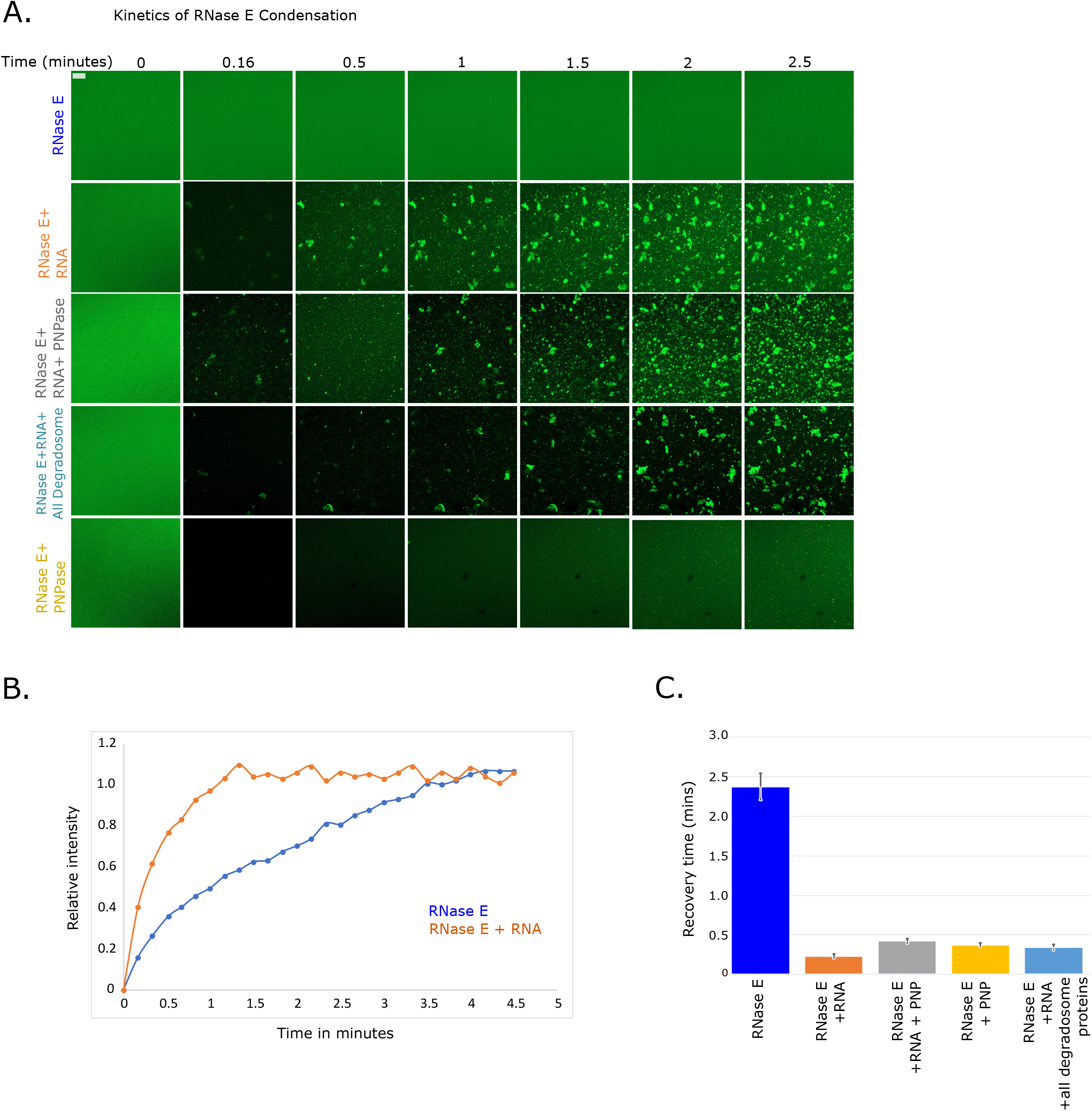
RNA stimulates rapid BR-body condensation *in vitro*. A) Time lapse imaging of RNase E CTD YFP (RCY) condensate formation. 6 µM RCY assembly was monitored either alone or with different combinations of 100ng/µL RNA, 5 µM PNPase, or all of the degradosome proteins (5 µM PNPase, 4 µM RhlB, 5 µM RNase D1, and 5 µM Aconitase) as indicated on the left. The images were captured at 20X magnification at the time points indicated above. The scale bar is 20 µm. B) Quantification of condensate growth by calculation of the % area occupied in the time lapse experiments from panel A. The data were quantified in ImageJ, averaged from three independent replicates, and plotted in MS Excel. C) Apparent FRAP recovery for droplets with the indicated composition. Because no RCY condensates were present at 6 µM, the concentration was increased to 30 µM to induce condensates. The values are averages from at least six different droplets, and the error bars represent the standard deviation.

RNase E is known to scaffold the RNA degradosome protein complex, and the degradosome proteins stoichiometrically bind to RNase E^31^ and are homogeneously present with RNase E in BR-bodies^4^. To test the effects of degradosome proteins on condensation kinetics, we added either PNPase or the full set of core degradosome proteins (PNPase, Aconitase, RhlB, and RNase D1) to the condensation assembly experiment. Upon the addition of PNPase or the full set of core RNA degradosome proteins, RNase E still rapidly condensed on a sub-minute timescale that was accelerated by the addition of RNA (Fig 7A). This suggests that the combined interactions with the RNA degradosome proteins likely do not dramatically alter RNase E’s condensation kinetics in the presence of RNA. This may be due to the degradosome protein binding sites being located outside the Arg-rich RNA binding sites in the C-terminal domain of RNase E^31^. As noted previously, we observed that the degradosome protein PNPase was able to stimulate RNase E condensation *in vitro* in the absence of RNA (Fig S5A). We therefore assayed the kinetics of RNase E and PNPase condensation in the absence of RNA and found that the kinetics were dramatically slower than in the presence of RNA (Fig 7A). Overall, this suggests that RNA provides the strongest kinetic stimulation of condensation, and the RNA degradosome clients can provide a slight further enhancement of the rate of RNA-mediated condensation.

To understand if RNA or degradosome protein interactions alter the mobility of molecules in the condensed state, we performed FRAP experiments (Fig 7B, C). RNase E condensates lacking RNA had the slowest recovery, at 2.4 min τ_1/2_, while the addition of RNA accelerated the FRAP recovery τ_1/2_ to 0.23 min. Interestingly, mobility was still fast upon the addition of PNPase and RNA or all of the degradosome proteins and RNA, with recovery times similar to the RNase E with RNA sample (τ_1/2_ of 0.34 and 0.42 min, respectively). This suggests that the addition of RNA significantly increases the mobility of RNase E condensates, regardless of the presence of degradosome clients. Interestingly, the addition of RNase E and PNPase in the absence of RNA showed a recovery τ_1/2_ of 0.38 min. This suggests that, even though the degradosome proteins do not dramatically stimulate the rate of condensation, degradosome protein interactions likely do influence the mobility of RNase E within the droplets, either indirectly by altering the structure of the CTD or directly by influencing multivalent protein-protein interactions in the condensed state. Altogether, RNA appears to stimulate the rate of RNase E condensation, while the recruitment of RNA or protein clients increases the mobility of RNase E within the condensate, which may promote enhanced biochemical activity.

## Discussion

### A complex protein interaction network impacts BR-body condensation and composition

Analysis of the BR-body proteome revealed enrichment of >100 proteins of various biochemical functions, suggesting that BR-bodies are more complex than previously assumed. We observed that a subset of BR-body enriched proteins known to stoichiometrically assemble into the RNA degradosome (RNase E, RhlB, Aconitase, and PNPase) appear to be uniformly enriched in BR-bodies *in vivo*^4^ and selectively recruited to RNase E condensates *in vitro* (Fig 6). This suggests that the RNA degradosome likely composes the “core” set of proteins making up each BR-body. As observed for yeast P-bodies, a set of “core” proteins that were found to be quantitatively enriched^27^ and likely homogeneously represented in these structures, were successfully reconstituted *in vitro* with similar dynamics properties and stoichiometries identified^28^. An important future goal will be to define the role of BR-body localization to each of the core enzymes involved in the RNA decay process. Indeed, in a two protein BR-body system using only RNase E and PNPase, PNPase activity was observed to be directly stimulated by condensation with RNase E^17^, suggesting that the organization in BR-bodies may help to stimulate RNA degradosome activity.

In addition to a uniformly enriched core of RNA degradosome proteins, we also found multiple proteins that were heterogeneously localized with BR-bodies, including the DEAD box RNA helicase RhlE, the transcription termination factor Rho, and the RNA chaperone Hfq. Although Rho was strongly colocalized with RNase E foci *in vivo*, we found that only a subset of RNase E foci contained Rho signal (Fig 2). RhlE and Hfq were both present in a subset of RNase E foci, and they could also form independent foci *in vivo* that did not contain RNase E (Fig 2). Importantly, non-RNase E scaffolds that can undergo heterotypic phase separation with RNA are enriched in BR-bodies, which suggests that many cellular RNP condensates exist and interact with BR-bodies. Five proteins that interact with RNase E were identified as scaffolds that can phase separate with RNA *in vitro*: RhlE, DEAD, Hfq, FabG, and MetK. Together with the RNApol and Rho condensates identified previously^5, 44^, this suggests that bacteria contain many RNP condensates that likely help coordinate multi-step reactions in RNA processing^3^.

DEAD box RNA helicases appear to be important players in biomolecular condensates^40, 45^, and we observed three enriched in BR-bodies. We observed that RhlE and DEAD act as scaffolds that undergo RNA-dependent phase separation *in vitro*, while RhlB cannot phase separate alone, but can enter the BR-body through RNase E’s IDR, similar to what was observed for their *E. coli* homologues^40^, suggesting phase separation is a conserved feature of these helicases. Interestingly, we observed strong recruitment of RhlE and RhlB into RNase E condensates, but DEAD was observed to be recruited to a lower extent (Fig 4). While RhlB was strongly colocalized with RNase E^4^ and RhlB foci dissociated in cells after RNase E depletion, RhlE showed heterogenous, partially overlapping localization with RNase E and remained localized in foci even after RNase E depletion (Fig 2). The function of DEAD box proteins in BR-bodies is not well understood; however, RhlE becomes more highly expressed when *C. crescentus* is grown in the cold^35^, which may alter the composition and function of BR-bodies in these conditions.

BR-body proteomics identified Hfq as a BR-body enriched protein. Hfq chaperones small RNAs to their target mRNAs and represses translation/induces decay^46–50^. It is unsurprising that Hfq was enriched in BR-bodies because the BR-body transcriptome is highly enriched with poorly translated mRNAs and small RNAs^29^. Hfq phase separates with RNA *in vitro* and forms heterogenous colocalized foci in *C. crescentus*. *E. coli* Hfq was recently found to phase separate with DNA, polyphosphate^41^, and RNA^10^, and its condensation is promoted *in vitro* by its intrinsically disordered CTD^41^. *E. coli* Hfq foci were previously identified to colocalize with RNase E in nitrogen starved *E. coli* cells^7^, and, while Hfq is not a core member of the *E. coli* or *C. crescentus* RNA degradosome, it has been found to associate with RNase E in multiple species^51^. Interestingly, we observed that *C. crescentus* Hfq is readily recruited into RNase E-RNA condensates, but RNase E was not recruited into Hfq-RNA condensates, suggesting a preferred directionality in forming a mixed condensate. *E. coli* Hfq condensates promote sRNA-mRNA annealing^10^ and RNase E IDR mutants prevent sRNA-mRNA silencing^52^. These RNase E IDR mutants were later found to prevent BR-body phase separation ^4^ Therefore, the coordination of Hfq with BR-bodies likely helps degrade translationally repressed mRNAs; the same function has been proposed for miRNA/siRNA/LSM in P-bodies^22^.

The BR-body associated proteins identified that can dissolve BR-bodies, ribosomal protein S1 and MetK, may play important regulatory roles in controlling BR-body assembly. Ribosomal protein S1 appears to dissolve RNase E condensates via its RNA binding capacity, while MetK appears to dissolve RNase E via protein-protein interaction with RNase E (Fig 5). Ribosomal protein S1 is present in a ribosome-bound and a ribosome free form; it unfolds structured mRNAs in preparation for translation initiation^53^ and protect mRNAs from RNase E-dependent decay^54^. Since BR-bodies and ribosomes compete for mRNA substrates^4^, ribosomal protein S1 may prevent BR-body condensation on highly translated mRNAs by locally disrupting RNase E condensation when in polysomes. Alternatively, if ribosomal protein S1 dissociates from the ribosome, it may be able to globally dissolve BR-bodies, potentially acting as a sensor of translation. We found that MetK, an essential enzyme that generates S-adenosylmethionine, could phase separate with RNA. MetK was previously found to bind RNase E in cells grown in the cold ^35^, and we identified a protein-protein interaction between RNase E and MetK under standard growth temperature (Fig 5, Fig S5C), suggesting MetK likely inhibits RNase E condensation directly by blocking multivalent interactions with RNase E or indirectly by altering the conformation of RNase E. Despite its rather weak RNA binding observed *in vitro*, *C. crescentus* MetK phase separated in the presence of RNA, and homologs from *S. meliloti* and *E. coli* were identified to bind RNA^55^. Because MetK is the sole enzyme creating S-adenosylmethionine in *C. crescentus*, perhaps it might act as a metabolic sensor that could tune the number of BR-bodies to carbon availability. In line with this hypothesis, MetK protein levels are dramatically altered upon carbon starvation^56^.

### RNA is critically important to BR-body condensation

Excitingly, reconstitution of minimal BR-body cores revealed that RNA, which was hypothesized to play a key role in BR-body assembly *in vivo*^4^, plays a key role in BR-body assembly *in vitro*. We observed that RNA can kinetically stimulate the rate of BR-body condensation on the sub-minute timescale (Fig 7), further suggesting that the main cause of RNase E’s robust foci dissolution under rifampicin treatment is the RNA depletion induced by rifampicin and not other cellular perturbations from this drug, such as nucleoid expansion^57^. The addition of RNA degradosome proteins does not alter the rate of RNase E condensation, but it does appear to enhance RNase E mobility by FRAP (Fig 7). *In vivo*, the RNA present in BR-bodies is likely mostly long, poorly translated mRNAs and small RNAs; BR-bodies are depleted of rRNA and other highly structured ncRNAs^29^. This suggests that, at least under logarithmic growth conditions, the availability of poorly translated RNA helps catalyze rapid BR-body assembly. Interestingly, BR-bodies could be induced with some client proteins in the absence of RNA, such as PNPase; however, the rate of the condensation was far slower than in the presence of RNA. While likely not relevant under log phase growth conditions as BR-bodies were observed to have sub-minute dynamics^4^, such assemblies might occur under non-growing conditions, like stationary phase.

The identification of five new RNP condensates associated with BR-bodies suggests these structures likely have a broader role in bacterial cell organization than previously anticipated. Indeed, recent high-throughput methods designed to detect proteins crosslinked to RNA identified >1100 RNA binding proteins in *E. coli*, including many metabolic enzymes^58, 59^. Similarly, we identified two metabolic enzymes, MetK and FabG, that have the capacity to phase separate with RNA (Fig 3). While the ability of metabolic enzymes to bind to RNA is referred to as “moonlighting” RNA binding activity^60^, it is possible that due to the large number of RNA binders, RNP condensates play a broad role in organizing metabolic pathways in the bacterial cytoplasm. Because experimentation to identify and characterize new condensates is a slow process, bioinformatic prediction of proteins that can undergo phase separation is the only method that can be utilized across whole proteomes. However, the current algorithms^61, 62^ do not accurately predict the phase separation of all five of the proteins we identified. Perhaps current phase separation prediction algorithms rely too much on IDR predictions, which often affect phase separation, but are not always required^63^. When developing next generation algorithms to predict phase separation, it will be important for the training data sets to include bacterial phase separating proteins to improve their predictive power.

## Materials and Methods

### Caulobacter crescentus cell growth

All *C. crescentus* strains used in this study were derived from the wild-type strain NA1000^64^ and were grown at 28^°^C in peptone-yeast extract (PYE) medium or M2 minimal medium supplemented with 0.2% D-glucose (M2G)^65^. When appropriate, the indicated concentration of vanillate (0.5 mM), xylose (0.2%), gentamycin (Gent) (0.5 mg/mL), kanamycin (Kan) (5 mg/mL), spectinomycin (Spec) (25 mg/mL), and/or streptomycin (Strp) (5 mg/mL) was added. Strains were analyzed at mid-exponential phase of growth (OD 0.3-0.6). Optical density was measured at 600 nm in a cuvette using a Nanodrop 2000C spectrophotometer.

### BR-body enrichment assay and LC-MS proteomics

BR-body enrichment was performed as in^30^. Enriched BR-bodies were then resuspended in buffered SDS before proteomics sample preparation.

#### Suspension Trapping (S-Trap) Sample Preparation

100 µg protein lysate (JS221 or JS299) was prepared in triplicate according to the manufacturer’s instructions. Trypsin was resuspended in a 50 mM ABC buffer and added to S-Traps at a 1:50 (enzyme:protein w/w) ratio and incubated for 8 hours at 37ºC. After the digestion, peptides were eluted according to the manufacturer’s protocol and vacuum centrifuged to dryness prior to desalting.

#### Solid Phase Extraction (SPE)

All samples were desalted prior to MS analysis with 10mg HLB cartridges according to manufacturer’s instructions as in ^66^. Samples were dried prior to analysis and stored at -80^°^C until re-suspension for LC-MS.

#### Analysis via Nano UHPLC-MS/MS

Dried digests were resuspended in 0.1% formic acid in water to a concentration of 250 ng/µL. 2 µL of solution was injected onto a 100mm × 100µm C_18_ BEH reverse phase chromatography column with an autosampler (nanoACQUITY, Waters Corporation). Peptides were separated over a 95 minute segmented gradient from 4-35% B (A= H_2_O + 0.1% FA, B=acetonitrile + 0.1% FA). MS-MS/MS was performed on a Q-Exactive HF mass spectrometer via electrospray ionization (Thermo Fisher Scientific). Samples were collected in biological triplicate, and data were acquired in technical triplicate injections using a TOP17 data-dependent method (DDA).

#### Peptide-Spectrum Matching (PSM)

RAW files from technical and biological triplicates were processed for peptide identification, protein inference, and false-discovery rate using the MetaMorpheus search engine^67^. A *C. crescentus* FASTA from UniProt (proteome ID UP000001364) was used with a common contaminant file. Data were first calibrated prior to search (‘Traditional’) using the default parameters. For ‘Main’ search, additional parameters included a maximum of two missed cleavages, two variable modifications per peptide, and a minimum peptide length of seven amino acids. The precursor mass tolerance was set to 5 ppm and the product mass tolerance was set to 20 ppm. The ‘common’ fixed and ‘common’ variable modifications were selected, and identifications were filtered to a q-value of <0.01. RAW data files are available through MassIVE with accession number (MSV000090894).

### *In vivo* localization of BR-body associated proteins

All *C. crescentus* protein fusion strains were grown in PYE-Kan or PYE-Gent media overnight. The next day, from the mid log-phase cultures, serial dilutions were done in M2G medium containing the appropriate amount of Kan or Gent and cells were grown overnight. The next day, the mid log-phase cultures were induced with 0.5 mM vanillate and grown for 6 hours. 1µL of the cells were spotted on a M2G 1.5% agarose pad, a cover slip was added, immersion oil was spotted on the cover slip, and the cells were imaged using a YFP filter cube.

### RNase E depletion

Depletion strains containing the xylose-inducible copy of RNase E were first grown overnight in M2G-Kan-Gent medium containing 0.2% xylose. The next day, from the log-phase cultures, serial dilutions were prepared in M2G-Kan-Gent medium containing 0.2% xylose overnight. Mid-log phase cells were then washed 3 times with 1 mL growth medium, resuspended in growth medium, and split equally into two tubes. 0.2% xylose was added to one tube. Depletion strains were analyzed at the mid-exponential phase of growth (OD 0.3-0.6) and after 24 hours of depletion of xylose. 1 µL of the cells under each condition were spotted on a M2G 1.5% agarose pad, a cover slip was added, immersion oil was spotted on the cover slip, and the cells were imaged using a YFP filter cube.

### Colocalization of YFP tagged foci forming proteins with BR-body markers

Dual-labeled strains expressing foci-forming proteins in Fig 2A with BR-body markers RNase E or Aconitase were first grown in PYE medium containing the appropriate amount of Kan, Gent, Spec or Strp overnight. The next day, from log-phase cultures, serial dilutions were done in M2G medium. The following day, mid log-phase cells were split into two tubes, and one was treated with an inducer (0.5 mM vanillate or 0.02% xylose), and both were grown for 6 hours. 1 µL of the cells under each condition was spotted on a M2G 1.5% agarose pad, a cover slip was added, immersion oil was spotted on the cover slip, and the cells were imaged using YFP, CFP, GFP, and TX-Red filter cubes. A dual-labeled strain expressing PopZ-mCherry with RNase E-GFP was used as a negative control. Fluorescence level analysis of the droplets was performed using MicrobeJ^68^. Line traces were performed using ImageJ and the correlation analysis between channels was performed in Microsoft Excel.

### *In vivo* localization of RNase E-GFP foci in the strains with BR-body enriched proteins deleted

JS671, JS672, JS735, and JS744 strains were grown in PYE medium containing the appropriate amount of Kan, Gent, Spec or Strp overnight. The next day, from the log-phase cultures, serial dilutions were done in M2G medium overnight. The following day, 1µL of the mid-log cells were spotted on a M2G 1.5% agarose pad, a cover slip was added, immersion oil was spotted on the cover slip, and the cells were imaged using a GFP filter cube.

### Dissolution of MetK and ribosomal protein S1

Strains expressing MetK and ribosomal protein S1 with BR-body marker Aconitase-mCherry were first grown overnight in PYE-Kan-Spec-Strp medium in 3 serial dilutions. Mid log-phase cells were split equally into two tubes, one was treated with 0.2% xylose, and the cells were grown for 6 hours. 1 µL of the cells under each condition was spotted on a M2G 1.5% agarose pad and imaged using TX-Red filter cubes.

### Cell imaging

Around 1 µL of cells were immobilized on 1.5% agarose pads made with M2G medium on microscope slides (Thermo Fisher Scientific 3051). Cell droplets were allowed to air dry, were covered by a cover slip, and imaged with immersion oil. All images were collected using a Nikon Eclipse NI-E with CoolSNAP MYO-CCD camera and 100x Oil CFI Plan Fluor (Nikon) objective, driven by Nikon elements software. The filter sets used for YFP, CFP, GFP, and mCherry imaging were chroma 96363, 96361, 96362, and 96322 models respectively. Cell image analysis was performed using MicrobeJ^68^.

### RNA extraction

For RNA extraction, mid-log phase cells (0.3 to 0.5 OD600) were pelleted at 20,000 g for 10 minutes in a microcentrifuge. Then, the cells were resuspended in 1mL 65^°^C Trizol (1mL for each 1 mL of NA1000 cells) and incubated at 65^°^C for 10 min in a heat block. 200 µL of chloroform was added and incubated for 5 min at room temperature. The samples were then spun at max speed (20,000 g) in a microcentrifuge for 10 min at room temperature. The aqueous layer was removed and RNA samples were precipitated using 700 µL isopropanol at -80^°^C for 1 hour and spun at 20,000 g for 1 hour at 4^°^C. The supernatant was removed and the pellet was washed with 1 mL of 70% ethanol. The samples were then spun again for 10 min at 20,000 g at 4^°^C, the supernatant was removed, and the pellet was resuspended in RNase-free water. RNA samples were run on 1X TBE/7M urea denaturing PAGE gels and visualized with SYBR Gold nucleic acid gel stain.

### Protein purifications

The protein coding genes were cloned into pET His6 MBP TEV,pET His6 G TEV, or pET28a plasmids and the expressed proteins were purified by His tag affinity purification, followed by the removal of the MBP or G tag by TEV proteolysis. RNase E CTD, RhlB, RhlE-IDR were expressed as N-terminal MBP fusions and all the other proteins were expressed as N-terminal G protein fusions. Briefly, the proteins were expressed in BL21 DE3 cells by inducing with 0.5mM IPTG at 37° C for 3.5 hours in 2X Luria-Bertani medium. The cells were then pelleted at 5000 rpm for 20 minutes and were lysed in a lysis buffer (20mM Tris pH 7.4, 500mM NaCl, 10% glycerol, 10mM imidazole, 1mM PMSF, 1 tablet protease inhibitor cocktail, and DNase I) at 4° C using a sonicator with 10 seconds pulse on and 30 seconds pulse off times for 18 cycles. The lysed cells were centrifuged at 15,000 rpm for 45 minutes and the resultant supernatant was loaded onto equilibrated Ni-NTA resin, the protein-bound resin was washed with 10 column volumes each of lysis buffer, chaperone buffer (20 mM Tris pH 7.4, 1 M NaCl, 10% glycerol, 10 mM imidazole, 5 mM KCl, 10 mM MgCl2, and 2 mM ATP), and low salt buffer (20 mM Tris pH 7.4, 1 M NaCl, 10% glycerol, and 10 mM imidazole). The proteins were eluted in elution buffer (20 mM Tris pH 7.4, 150 mM NaCl, 10% glycerol, and 250 mM imidazole) and dialyzed into storage buffer (20 mM Tris pH 7.4, 150 mM NaCl). The contaminant proteins were removed by passing through a S200 size exclusion column. The concentrated proteins were stored at -80°C.

RNase E CTD purification: His6-MBP-TEV-RNase E CTD was expressed and purified in the same way as described above except the lysis buffer additionally contained 1M NaCl and 0.1% Triton X-100.

Hfq purification: Hfq protein was expressed at 18°C for 16 hours. The lysis buffer and elution buffer contained 1 M NaCl instead of 150 mM and 500 mM NaCl, respectively. To remove the contaminating nucleic acids, the His tag affinity purified protein was treated with 30 µg/mL RNase A and 5 U/mL DNase I for 1 hour at 37° C, concentrated, and loaded onto a HiLoad 16/60 Superdex 200 size exclusion column (GE healthcare). The absence of RNase A was further validated by incubating RNA with the protein.

### Cyanine 5 Protein labeling

Labeling reactions were carried out according to the manufacturer’s protocol (Lumiprobe). Cy5 NHS ester dye dissolved in DMSO was incubated with proteins at an equal ratio in 0.1M sodium bicarbonate buffer pH 8.5 at room temperature for 4 hours. A protein labeling calculator (https://www.lumiprobe.com/calc/protein-labeling) was used to calculate the amount of protein and dye used in the labeling reactions. The excess dye was removed by dialyzing against protein storage buffer (20mM Tris pH 7.4, 150mM NaCl) using a 3 kDa cut off dialysis cassette. Degree of labeling was calculated using the formula:

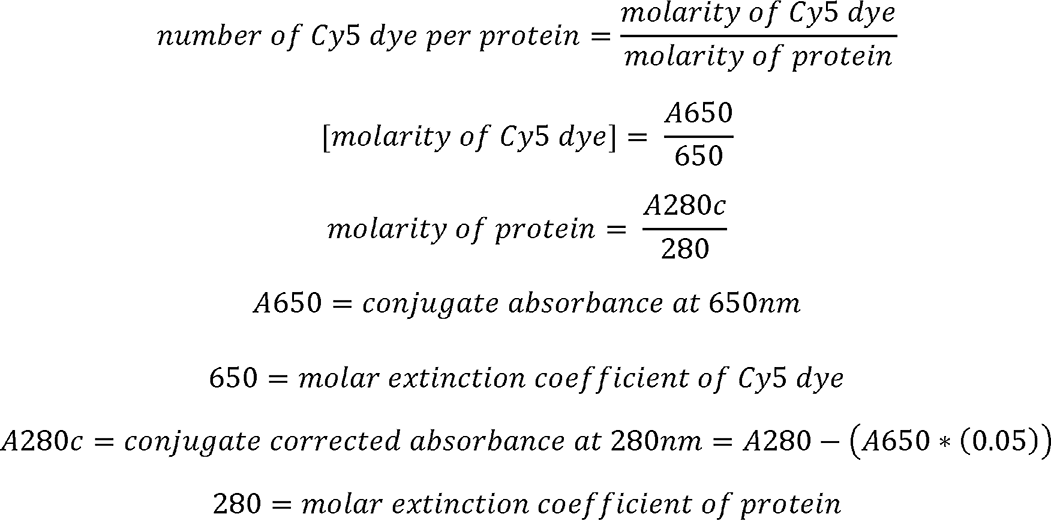

### phase separation experiments and droplet assays

Proteins were incubated at their *in vivo* concentrations in the presence or absence of *C. crescentus* total RNA (20 ng/µL) in 20mM Tris pH 7.4, 100mM NaCl for 30 minutes at room temperature. The mixture was pipetted onto a glass slide, covered with a cover slip, and imaged using a Nikon epifluorescence microscope. For quantification of the number and distribution of droplets, 10 images for each protein were analyzed using the “analyze particles” function in ImageJ..

For the recruitment assays, 6 µM RNAse E CTD was incubated with 20 ng/µL of *C. crescentus* total RNA in 20mM Tris pH 7.4, 100mM NaCl for 20 minutes at room temperature, followed by the addition of Cy5-labeled client proteins and an additional incubation for 20 minutes before imaging. The level of recruitment was assessed by calculating the partition coefficient of client proteins. 30 droplets were analyzed in each case.

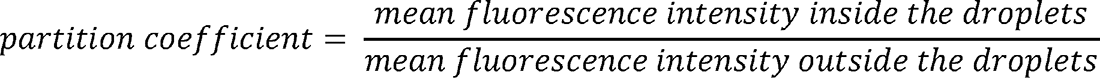

ImageJ software was used to quantify the mean fluorescence and the data were plotted in PlotsOfData^69^. Co-assembly experiments were performed by mixing all the components and incubating for 30 minutes at room temperature before imaging.

For the salt and RNA dependency experiments, preformed protein and RNA condensates were incubated with either varying concentrations of NaCl or RNase A for 30 minutes at room temperature before the solution was imaged on a glass slide using phase contrast microscopy.

RNase E condensation induction in the absence of RNA was carried out by adding client proteins at varied concentrations to 6 µM RNase E CTD in 20mM Tris, 100mM NaCl and incubating for 30 minutes at room temperature. The solution was imaged on a glass slide using phase contrast microscopy.

### *In-vitro* protein pull down assay

The RNAse E CTD tagged with maltose binding protein (MBP) or MBP only was incubated with bait proteins in pull down binding buffer (20mM HEPES pH 7.4, 100mM NaCl, 2% glycerol, and 2mM DTT) for 60 minutes at room temperature. 100 µL of dextrin agarose resin was added to the above mixture and incubated for another 90 minutes at 4°C on a tube rotator. The samples were washed four times with 1 mL of wash buffer (20mM HEPES pH 7.4, 100mM NaCl). Bound proteins were eluted with 100 µL of elution buffer (20mM HEPES pH 7.4, 150mM NaCl, 10mM maltose) at 30°C with constant shaking. The eluted proteins were resolved on an SDS PAGE gel.

### Electrophoretic mobility shift assay

RNA (50 nM) was incubated with the specified concentrations of proteins for 30 minutes at room temperature in binding buffer (20mM Tris pH 7.4, 150mM NaCl, 5% glycerol, 2mM DTT, 1mM EDTA, and 10 µg/mL BSA). The incubated samples were resolved on a 5% acrylamide/bis-acrylamide native gel (prepared in TBE buffer) at 4°C in 0.5X TBE running buffer. The gel was stained with SYBR Gold for 20 minutes and scanned using a Typhoon scanner.

### Bacterial Two-Hybrid assay

The bacterial two-hybrid assay was carried out as described previously in^42^. The bait protein RNase E was cloned into the pKNT25 plasmid (kanamycin resistance) using Xba1 and EcoR1 sites and the prey protein MetK was cloned into the pUT18 plasmid (ampicillin resistance) using Xba1 and Sac1 sites. The sequence verified plasmids were transformed into chemically competent BTH101 cells (adenylate cyclase deficient) and the colonies were selected on LB-agar plates containing kanamycin (50 µg/mL), ampicillin (100 µg/mL), X-gal (40 µg/mL), and IPTG (0.5mM), and incubated at 30°C for 72 hours. The resultant blue colonies were grown in LB-medium containing kanamycin (30 µg/mL) and ampicillin (50 µg/mL) and log phase cells were plated onto LB-agar plates containing kanamycin, ampicillin, X-gal, and IPTG as mentioned above. The positive control cells were transformed with pKNT25-Zip and pUT18-Zip plasmids, which upon expression reconstitute a functional adenylate cyclase enzyme. The negative control cells were transformed with the pKNT25 and pUT18 plasmids.

### Determination of the assembly order of RhIE, RNase E, and Aconitase foci

*C. crescentus* JS716 (RhIE-YFP/Aconitase-mCherry) and JS452 (RNase E-YFP/Aconitase-mCherry) were grown overnight in PYE medium containing kanamycin (5 μg/mL), spectomycin (25 μg/mL), and streptomycin (5 μg/mL) until the cultures reached mid log-phase (OD ∼0.3). Prior to imaging, the cells were diluted to 0.05 OD in PYE media and induced with 0.02% xylose for 15 minutes at 28°C. Aliquots of 1.5 μL of culture were spotted onto pads of 2% agarose in M2G medium and sandwiched between two coverslips. Cells were imaged with an Olympus IX71 inverted epifluorescence microscope with a 100× objective (NA 1.40, oil immersion) kept at 28 °C using an objective heater (Bioptechs). The YFP fusion proteins were imaged under excitation from a 488-nm laser (Coherent Cube 488-50; 0.4 W cm^−2^) and the mCherry fusions were imaged under excitation from a 561-nm laser (Coherent Sapphire 561-50; 2.0 W cm^−2^). Both channels were imaged simultaneously on a 512 × 512 pixel Photometric Evolve electron-multiplying charge-coupled device (EMCCD) camera, using an OptoSplit II image splitter (Cairn Research) with a 635-nm long pass filter and a 525/50 nm bandpass filter. An integration time of 250 ms with a delay between frames of 5 s was used, where a shutter blocked irradiation during the delay time to minimize photobleaching of YFP and mCherry.

To analyze the fluorescence intensities, the mean intensity of each cell in each channel was first subtracted from the movies. Foci were then detected by eye in the background-subtracted movies, and the mean and maximum intensities within the 9 × 9 pixel region of interest (ROI) surrounding each focus was measured in both channels. Time zero was set to be the frame immediately preceding the first frame with an ROI mean intensity at least 10% higher than the maximum intensity value in the ROI in the previous frame. This time zero was also assigned as time zero on the other channel. Subsequently, the maximum intensity within the ROI was divided by the maximum intensity of the ROI in the time zero frame to determine the fluorescence intensity enhancement of each focus at each time.

### Condensate growth kinetic assays

Time lapse images were acquired using a Zeiss LSM 800 microscope. 6 µM of RNase E CTD YFP (RCY) was spotted onto a glass slide, and either RNA or proteins or both at the specified concentrations were pipetted onto the RCY solution. Image acquisition was started before the addition of RNA or proteins or both, but the zero minute time point was calculated from the time they were added to the RCY solution on the glass slide. The zero minute time point for ‘RCY only’ started from the time the sample was spotted onto the glass slide. The images were acquired for 2.5 minutes with a 50 ms time interval. The data were quantified in ImageJ and plotted in MS Excel.

## QUANTIFICATION AND STATISTICAL ANALYSIS

### Foci quantification

*In vivo* protein fusion foci were quantified using MicrobeJ^68^. Cell outlines were first identified using the medial axis algorithm in the phase channel with a minimal cell length of 1.35 µm and the segmentation option. Cell outlines were then manually curated to remove cells containing erroneous outlines. Next, the maxima function was used with the ‘‘foci’’ option with a minimal area of 0.01 µm^2^ and minimal length of 0.1 µm with the segmentation option on and the association inside option on. Tolerance and Z-score parameters were tuned for each protein fusion type.

To quantify the average foci per cell of each strain, three replicates were performed on three different days and three images were analyzed from each day. We summed all foci identified in each cell and divided by the total number of cells to calculate number of foci per cell.

### Colocalization analysis

Dual channel images of enriched BR-body protein fusions with BR-body markers were analyzed via the line trace method to assess colocalization. For each focus identified, we drew a line through the focus and quantified the intensity vector across the line. We then analyzed the other channel across the same line, and calculated the correlation of the intensity of each channel. The images were analyzed for a minimum of 30 foci identified in the BR-body enriched protein (presented in Fig 2B). PopZ-mCherry was used as a negative control for the analysis.

### FRAP assays

FRAP was performed according to a standard protocol ^70^. Briefly, the measured F(t) over the bleached circular area of diameter (d) = 2 μm, normalized by that of the unbleached region of the same diameter, F(0), was fitted by a one-phase exponential function using the “Bottom to (Span+ Bottom)” analysis as F(t)/F(0) = Bottom + Span*(1 − exp(−t ln 2/τ_1/2_)), where τ_1/2_ is the halftime for diffusion. The apparent diffusion coefficient, D, was obtained from τ_1/2_ according to D =d2/(4τ_1/2_).

## Authors Contribution

VN performed protein purifications, all *in vitro* experiments, bacterial 2 hybrid assay, and *in vivo* experiments. IWR performed protein purifications, *in vivo* expression, depletion, and colocalization experiments, and made plasmids and strains for *in vivo* experiments. VN, AH, JV performed droplet kinetic assays and FRAP experiments. KM, AG, and YH purified proteins. NM purified BR bodies for BR-body proteomics. CBM and MC performed LC-MS data collection and analysis. LAOR performed RhlE *in vivo* assembly experiments. VN, IWR, and JMS wrote the paper.

## Acknowledgements

NIH grant R35GM124733 to JS. NIH grants R01GM143182 and R01GM144731 to JB. NIH Grant R01GM136863 to WSC. NSF grant CMMI-1914436 to YZ. NIH grant R01GM139277 to MC. We would like to acknowledge the Mass Spectrometry and Proteomics Facility at Notre Dame. *Caulobacter* cell and protein icons in the graphical abstract were created with Biorender.com. The authors thank Vincent Lawal for help with Bacterial 2 hybrid assay. The authors thank Erin Schrader for critical feedback.

