## Supplemental materials and methods for "The BR-body proteome contains a complex network of protein-protein and protein-RNA interactions"

**Supplemental Information**

**Supplemental Figure Legends**

**Figure S1**. **Subcellular localization patterns in *Caulobacter crescentus* of the YFP fusions.**

(A) Localization pattern of foci-forming YFP fusions. (B) Localization pattern of no foci-forming YFP fusions. (C) FabG YFP, RNaseT2 YFP, RNase L YFP, and RNase L1 YFP proteins were expressed at levels that were too low to measure foci. All genes were placed under control of the vanillate promoter grown in M2G minimal media. *In vivo* protein fusion strains were analyzed at mid-exponential phase of growth (OD 0.3-0.6) following 6-hours of induction with 0.5mM vanillate. 1 µL of the cells were spotted on a M2G 1.5% agarose pad, a cover slip was added, immersion oil was spotted on the cover slip, and imaged using a YFP filter cube.

**Figure S2**. Subcellular localization patterns of *C. crescentus* protein-YFP strains expressed from the *vanA* locus in the RNase E depletion background where the sole copy of the RNase E gene is controlled by the xylose promoter. The YFP intensity of aconitase-YFP was too low to measure when expressed from the vanillate promoter, so a native gene fusion was used instead. YFP intensity of each image was normalized from its brightest level relative to its background level. Depletion strains were analyzed at mid-exponential phase of growth (OD 0.3-0.6) and after 24 hours of depletion of xylose. Three replicates were performed in three different days. 1 µL of the cells under each condition were spotted on a M2G 1.5% agarose pad, a cover slip was added, immersion oil was spotted on the cover slip, and imaged using a YFP filter cube. Scale bar is 3 µm for all images.

**Figure S3**. (A) Purified proteins on the SDS PAGE gel. Proteins from 1 to 18 were resolved on 10% gel and the proteins from 19 to 22 were resolved on 15% gel. 1. RNase E CTD (49.4 kDa) 2. PNPase (76.4 kDa) 3. Aconitase (96.6 kDa) 4. MBP-RhlB (99 kDa) 5. RNase D (42.8 kDa) 6. G-RhlE (57.3 kDa) 7. G-DeaD full length (87 kDa) 8. DnaK (48.6 kDa) 9. FabG-6X His (25.3 kDa) 10. NudC-6XHis (35 kDa) 11. RppH-6XHis (20.6 kDa) 12. Rho (53.3 kDa) 13. G- NusG (27.5 kDa) 14. G-Tryosyl tRNA ligase (52 kDa) 15. SpmX-6X His (46.8 kDa) 16. Ribosomal protein S1-6X His (63 kDa) 17. metK-6X His (44.6 kDa) 18. RNase E CTD eYFP-6X His (80.5 kDa) 19. Hfq full length-6X His (10.2 kDa) 20. G-Hfq (1 – 66)-6X His (14.5 kDa) 21. RhlE (1-360) (39 kDa) 22. G-DeaD (1 – 520)-6X His (64 kDa) (B) Phase contrast microscope images of purified proteins that did not undergo phase separation. PNPase (23 µM), Aconitase (15 µM), RhlB (4 µM), RNase D (5 µM), NudC (2.5 µM), RppH (1 µM), Rho (5 µM), NusG (5 µM), DnaK (5 µM), Tryrosyl tRNA ligase (5 µM), S1 protein (50 µM) (scale is 5 µm for all images). (C) Phase contrast microscopy images of RhlE full length (2 µM), RhlE 1-360 (2 µM) Hfq full length (5 µM), Hfq 1-66 (5 µM), DeaD full length (5 µM), DeaD 1-520 (5 µM). Quantification of the droplets was performed in ImageJ and the data was plotted in PlotsOfData (scale is 5 µm for all images).

**Figure S4**. Recruitment of client proteins into RNase E condensates *in-vitro.* (A) Microscope images of the client proteins recruitment into RNase E condensates. RNase E CTD was unlabelled and the client protein was labeled with Cy5 dye (scale is 3 µm). (B) Partition coefficient of RNase E CTD Alexa Flour 350 in the presence of client proteins. Partition coefficient was quantified in ImageJ and the data plotted in PlotsOfData. 30 droplets were analyzed in each case and each dot in the graph represents a droplet.

**Figure S5**. (A) Phase contrast microscope images of RNase E CTD condensation by client proteins in the absence of RNA (scale is 5 µm). (B) RNase E co-precipitates with Aconitase and MetK. Left gel contains input controls and right gel contains pull down samples. Pulldown of 5 µM MBP-RNase E CTD or 5 µM MBP incubated with 5 µM of Aconitase, 15 µM MetK and 15 µM S1 . Eluates were run on 10 % SDS-PAGE and stained with SimplyBlue SafeStain (Thermo Fisher). MBP mock reactions were used as a negative control. (C) Recruitment of scaffold proteins by scaffold protein condensates in the absence of RNA. Scaffold protein condensation was stimulated by increasing its concentration in the absence of RNA followed by the addition of a Cy5 labeled scaffold protein at the in-vivo concentration, and the partition coefficient was calculated. In the table given are the partition coefficient values.

**Figure S6.** (A) Single molecule analysis of RhlE YFP and Aconitase mCherry foci assembly in the *C. crescentus* cells. (B) FRAP analysis of RNase E CTD YFP + total RNA and RhlE Cy5 + total RNA condensates *in-vitro.* The plot is an average of three condensates and the error bars represent standard deviation.

**Figure S7.** (Top) 6 µM RNase E for 30 mins incubation. (Bottom) 20 µM RNase E for 30 mins incubation.

**Strain and Plasmid Construction**

All strains are listed in Table S3, plasmids in Table S4, and DNA oligos in Table S5. A detailed protocol for the generation of all plasmids and strain are listed below. In all cases, YFP listed in eYFP from Thanbichler *et al.* NAR 2007.

**Plasmid Construction**

***Protein expression plasmids***

**NudC-pET28a-2** plasmid was generated by subcloning from pVNudC-YFPC-2 into pET28a-2 plasmid. First, both pVNudC-YFPC-2 and pET28a-2 were cut with Nde1/EcoRI enzymes. Next, the digested fragments were gel purified, then ligated using T4 DNA ligase enzyme. The ligation was transformed into chemically competent E. coli cells and selected on LB-Kan plates. The resulting kanR colonies were then screened by PCR for the insert and verified by sanger sequencing (genewiz).

**RppH-pET28a-2** plasmid was generated by subcloning from pVRppH-YFPC-2 into pET28a-2 plasmid. First, both pVRppH-YFPC-2 and pET28a-2 were cut with Nde1/EcoRI enzymes. Next, the digested fragments were gel purified, then ligated using T4 DNA ligase enzyme. The ligation was transformed into chemically competent E. coli cells and selected on LB-Kan plates. The resulting kanR colonies were then screened by PCR for the insert and verified by sanger sequencing (genewiz).

**Rho-pET28a-2** plasmid was generated by subcloning from pVRhO-YFPC-2 into pET28a-2 plasmid. First, both pVRhO-YFPC-2 and pET28a-2 were cut with Nde1/EcoRI enzymes. Next, the digested fragments were gel purified, then ligated using T4 DNA ligase enzyme. The ligation was transformed into chemically competent E. coli cells and selected on LB-Kan plates. The resulting kanR colonies were then screened by PCR for the insert and verified by sanger sequencing (genewiz).

**MetK-pET28a-2** plasmid was generated by subcloning from pVMetK-YFPC-2 into pET28a-2 plasmid. First, both pVMetK-YFPC-2 and pET28a-2 were cut with Nde1/EcoRI enzymes. Next, the digested fragments were gel purified, then ligated using T4 DNA ligase enzyme. The ligation was transformed into chemically competent E. coli cells and selected on LB-Kan plates. The resulting kanR colonies were then screened by PCR for the insert and verified by sanger sequencing (genewiz).

**FabG-pET28a-2** plasmid was generated by subcloning from pVFabG-YFPC-2 into pET28a-2 plasmid. First, both pVFabG-YFPC-2 and pET28a-2 were cut with Nde1/EcoRI enzymes. Next, the digested fragments were gel purified, then ligated using T4 DNA ligase enzyme. The ligation was transformed into chemically competent E. coli cells and selected on LB-Kan plates. The resulting kanR colonies were then screened by PCR for the insert and verified by sanger sequencing (genewiz).

**HFQ-pET-28a(+)** plasmid was generated by GenScript as a gblock fragment. The gbock fragment was cloned into pET-28a vector by GenScript.

*Hfq gblock*

GGCCGCAAGCTTGTCGACGGAGCTCGAATTCgcgtcgtcggcgtcggcgctcggctcatagagctgaacgggttgagccggcatgatggtggaaatggcgtgcttgtagaccagttgcgactggccgtcgcgacgcagcaggacgcagaaattgtcgaaccagctgaccacgccctgcagcttgacgccattgacgaggaagatggtgagcggcgtcttcgacttgcgaacgctgttcaggaaggtgtcctgaagattttgcttcttttcggcggaCATATGGCTGCCGCGCGGCACCAGGCCGCTG

The protein expression genes in **pVN001**, **pVN004**, **pVN028**, **pVN030** plasmids were amplified from *C. crescentus* genomic DNA using VN003F & VN003R, VN001F & VN002R, VN004F & VN034R, VN035F & VN036R oligos respectively and cloned separately into pET-MBP-TEV-His vector using Ssp1 and HindIII restriction sites. The genes in **pVN008**, **pVN024**, **pVN027**, **pVN032**, **pVN034**, **pVN035, pVN025, pVN045** plasmids were amplified from *C. crescentus* genomic DNA using VN004F & VN004R, VN019F & VN020R, VN026F & VN027R, VN037F & VN037R, VN039F & VN039R, VN040F & VN040R, VN019F & VN021R, VN062F & VN063R oligos respectively and cloned separately into pET-MBP-TEV-His vector using Ssp1 and HindIII restriction sites.

**pMJC0100** plasmid was generated by amplifying the C. crescentus aconitase gene using primers MJC229 and MJC230 and inserted into pTEV5 using Gibson assembly. The Gibson reaction was transformed into chemically competent E. coli cells and selected on LB-Amp plates. The resulting ampR colonies were then screened by PCR for the insert and verified by sanger sequencing.

1) MJC229 CCAACTAGTGAAAACCTGTATTTTCAGGGCGCTATGGCGTCTGTGGACAGCCT
2) MJC230 AGAATTCCATGGCCATATGGCTTTAGTCGGCCTTGGCCAGGTTCCGCAGC

***Caulobacter plasmid designs***

All *Caulobacter crescentus* gene inserts were amplified by PCR from the NA1000 genome.

**PAPI-YFP** fusion was generated by cloning the Papl gene (CCNA_00413) lacking a stop codon in frame with YFP in the pVYFPC-2 plasmid^1^. First, the Papl gene was amplified by PCR from the NA1000 genome using the following DNA primers:

1) F_00413	cgaggaaacgcatatgagcagtgaaatgatcggcaacgcg

2) R_00413	cgttcgaattcgcgtagaccatgccctgggcgacg

Next, the resulting PCR product was digested with NdeI and EcoRI enzymes, gel purified, then ligated to NdeI/EcoRI cut pVYFPC-2 plasmid using T4 DNA ligase enzyme. The ligation was transformed into chemically competent E. coli cells and selected on LB-Kan plates. The resulting kanR colonies were then screened by PCR for the insert and verified by sanger sequencing (genewiz).

**RhlE-YFP** fusion was generated by cloning the RhlE gene (CCNA_00878) lacking a stop codon in frame with YFP in the pVYFPC-2 plasmid^1^. First, the RhlE gene was amplified by PCR from the NA1000 genome using the following DNA primers:

1) F_00878	cgatgcgaggaaacgcatatgactcaattttccgaccttggcct

2) R_00878	cgttcgaattcgcgtcgataggcgaccagcgc

Next, the resulting PCR product was digested with NdeI and EcoRI enzymes, gel purified, then ligated to NdeI/EcoRI cut pVYFPC-2 plasmid using T4 DNA ligase enzyme. The ligation was transformed into chemically competent E. coli cells and selected on LB-Kan plates. The resulting kanR colonies were then screened by PCR for the insert and verified by sanger sequencing (genewiz).

**Rho-YFP** fusion was generated by cloning the Rho gene (CCNA_03876) lacking a stop codon in frame with YFP in the pVYFPC-2 plasmid^1^. First, the Rho gene was amplified by PCR from the NA1000 genome using the following DNA primers:

1) F_03876	cgaggaaacgcatatgaccgaagacaccgaaaaccagg

2) R_03876	cgttcgaattcgcggtgttcatcgactggaagaagtc

Next, the resulting PCR product was digested with NdeI and EcoRI enzymes, gel purified, then ligated to NdeI/EcoRI cut pVYFPC-2 plasmid using T4 DNA ligase enzyme. The ligation was transformed into chemically competent E. coli cells and selected on LB-Kan plates. The resulting kanR colonies were then screened by PCR for the insert and verified by sanger sequencing (genewiz).

**RppH-YFP** fusion was generated by cloning the RppH gene (CCNA_03553) lacking a stop codon in frame with YFP in the pVYFPC-2 plasmid^1^. First, the RppH gene was amplified by PCR from the NA1000 genome using the following DNA primers:

1) F_03553	cgaggaaacgcatatgactgagctggaccatcctcaacatcg

2) R_03553	cgttcgaattcgcgttctccccctcgcggcg

Next, the resulting PCR product was digested with NdeI and EcoRI enzymes, gel purified, then ligated to NdeI/EcoRI cut pVYFPC-2 plasmid using T4 DNA ligase enzyme. The ligation was transformed into chemically competent E. coli cells and selected on LB-Kan plates. The resulting kanR colonies were then screened by PCR for the insert and verified by sanger sequencing (genewiz).

**NudC-YFP** fusion was generated by cloning the NudC gene (CCNA_00267) lacking a stop codon in frame with YFP in the pVYFPC-2 plasmid^1^. First, the NudC gene was amplified by PCR from the NA1000 genome using the following DNA primers:

1) F_00267	gcgaggaaacgcatatgcctctttcgatcatcaccaacacc

2) R_00267	cgttcgaattcgccgcctcttcggcccaggc

Next, the resulting PCR product was digested with NdeI and EcoRI enzymes, gel purified, then ligated to NdeI/EcoRI cut pVYFPC-2 plasmid using T4 DNA ligase enzyme. The ligation was transformed into chemically competent E. coli cells and selected on LB-Kan plates. The resulting kanR colonies were then screened by PCR for the insert and verified by sanger sequencing (genewiz).

**PNP-YFP** fusion was generated by cloning the PNPase gene (CCNA_00033) lacking a stop codon in frame with YFP in the pVYFPC-2 plasmid^1^. First, the PNPase gene was amplified by PCR from the NA1000 genome using the following DNA primers:

1) F_00033	cgatgcgaggaaacgcatatgttcgatatcaaacgcaagacgatcgagtggg

2) R_00033	cgttcgaattcgccgcctcttcggccgccgc

Next, the resulting PCR product was digested with NdeI and EcoRI enzymes, gel purified, then ligated to NdeI/EcoRI cut pVYFPC-2 plasmid using T4 DNA ligase enzyme. The ligation was transformed into chemically competent E. coli cells and selected on LB-Kan plates. The resulting kanR colonies were then screened by PCR for the insert and verified by sanger sequencing (genewiz).

**RNaseD1-YFP** fusion was generated by cloning the RNaseD1 gene (CCNA_01776) lacking a stop codon in frame with YFP in the pVYFPC-2 plasmid^1^. First, the RNaseD1 gene was amplified by PCR from the NA1000 genome using the following DNA primers:

1) F_01776	cgaggaaacgcatatgaagctgatcaccaccaccgcc

2) R_01776	cgtaacgttcgaattcgcatcgttcttgggggcgcg

Next, the resulting PCR product was digested with NdeI and EcoRI enzymes, gel purified, then ligated to NdeI/EcoRI cut pVYFPC-2 plasmid using T4 DNA ligase enzyme. The ligation was transformed into chemically competent E. coli cells and selected on LB-Kan plates. The resulting kanR colonies were then screened by PCR for the insert and verified by sanger sequencing (genewiz).

**RNaseD2-YFP** fusion was generated by cloning the RNaseD2 gene (CCNA_03717) lacking a stop codon in frame with YFP in the pVYFPC-2 plasmid^1^. First, the RNaseD2 gene was amplified by PCR from the NA1000 genome using the following DNA primers:

1) F_03717	gcgaggaaacgcatatggccaatttcgttcacgagggcg

2) R_03717	cgttcgaattcgcgctgtgggcgaagatgtccatctcc

Next, the resulting PCR product was digested with NdeI and EcoRI enzymes, gel purified, then ligated to NdeI/EcoRI cut pVYFPC-2 plasmid using T4 DNA ligase enzyme. The ligation was transformed into chemically competent E. coli cells and selected on LB-Kan plates. The resulting kanR colonies were then screened by PCR for the insert and verified by sanger sequencing (genewiz).

**RhlB-YFP** fusion was generated by cloning the RhlB gene (CCNA_01923) lacking a stop codon in frame with YFP in the pVYFPC-2 plasmid^1^. First, the RhlB gene was amplified by PCR from the NA1000 genome using the following DNA primers:

1) F_01923	acgatgcgaggaaacgcatatgactgaattcaccgacctagggctatcg

2) R_01923	atcttaaggtacccttcgcgccgcgcggcgg

Next, the resulting PCR product was digested with NdeI and EcoRI enzymes, gel purified, then ligated to NdeI/EcoRI cut pVYFPC-2 plasmid using T4 DNA ligase enzyme. The ligation was transformed into chemically competent E. coli cells and selected on LB-Kan plates. The resulting kanR colonies were then screened by PCR for the insert and verified by sanger sequencing (genewiz).

**AconA-YFP** fusion was generated by cloning the Aconitase gene (CCNA_03781) lacking a stop codon in frame with YFP in the pVYFPC-2 plasmid^1^. First, the Aconitase gene was amplified by PCR from the NA1000 genome using the following DNA primers:

1) F_03781	cgaggaaacgcatatggcgtctgtggacagc

2) R_03781	cgagatcttaaggtaccgtcggccttggccaggttc

Next, the resulting PCR product was digested with NdeI and EcoRI enzymes, gel purified, then ligated to NdeI/EcoRI cut pVYFPC-2 plasmid using T4 DNA ligase enzyme. The ligation was transformed into chemically competent E. coli cells and selected on LB-Kan plates. The resulting kanR colonies were then screened by PCR for the insert and verified by sanger sequencing (genewiz).

**RNaseJ-YFP** fusion was generated by cloning the RNaseJ gene (CCNA_02012) lacking a stop codon in frame with YFP in the pVYFPC-2 plasmid^1^. First, the RNaseJ gene was amplified by PCR from the NA1000 genome using the following DNA primers:

1) F_02012	tgcgaggaaacgcatatgaaaaagtccaagaacgacgag

2) R_02012	aacgttcgaattcgcaatgcgaagaaccgtgg

Next, the resulting PCR product was digested with NdeI and EcoRI enzymes, gel purified, then ligated to NdeI/EcoRI cut pVYFPC-2 plasmid using T4 DNA ligase enzyme. The ligation was transformed into chemically competent E. coli cells and selected on LB-Kan plates. The resulting kanR colonies were then screened by PCR for the insert and verified by sanger sequencing (genewiz).

**RNaseHI-YFP** fusion was generated by cloning the RNaseHI gene (CCNA_03476) lacking a stop codon in frame with YFP in the pVYFPC-2 plasmid^1^. First, the RNaseHI gene was amplified by PCR from the NA1000 genome using the following DNA primers:

1) F_03476	cgaggaaacgcatatgacgccgaaggtcacgatctataccg

2) R_03476	cgtaacgttcgaattcgcgatgacgcgcggattgg

Next, the resulting PCR product was digested with NdeI and EcoRI enzymes, gel purified, then ligated to NdeI/EcoRI cut pVYFPC-2 plasmid using T4 DNA ligase enzyme. The ligation was transformed into chemically competent E. coli cells and selected on LB-Kan plates. The resulting kanR colonies were then screened by PCR for the insert and verified by sanger sequencing (genewiz).

**RNaseHII-YFP** fusion was generated by cloning the RNaseHIl gene (CCNA_00383) lacking a stop codon in frame with YFP in the pVYFPC-2 plasmid^1^. First, the RNaseHII gene was amplified by PCR from the NA1000 genome using the following DNA primers:

1) F_00383	cgaggaaacgcatatgccgcccggacccgac

2) R_00383	gcgtaacgttcgaattcgcaaggtctagctcgccgttgacc

Next, the resulting PCR product was digested with NdeI and EcoRI enzymes, gel purified, then ligated to NdeI/EcoRI cut pVYFPC-2 plasmid using T4 DNA ligase enzyme. The ligation was transformed into chemically competent E. coli cells and selected on LB-Kan plates. The resulting kanR colonies were then screened by PCR for the insert and verified by sanger sequencing (genewiz).

**SmpB-YFP** fusion was generated by cloning the SmpB gene (CCNA_01254) lacking a stop codon in frame with YFP in the pVYFPC-2 plasmid^1^. First, the SmpB gene was amplified by PCR from the NA1000 genome using the following DNA primers:

1) F_01254	cgaggaaacgcatatgatgtccaagccgatcgcg

2) R_01254	cgtaacgttcgaattcgtcgccgcgatcgcccttc

Next, the resulting PCR product was digested with NdeI and EcoRI enzymes, gel purified, then ligated to NdeI/EcoRI cut pVYFPC-2 plasmid using T4 DNA ligase enzyme. The ligation was transformed into chemically competent E. coli cells and selected on LB-Kan plates. The resulting kanR colonies were then screened by PCR for the insert and verified by sanger sequencing (genewiz).

**RNaseL-YFP** fusion was generated by cloning the RNaseL gene (CCNA_01143) lacking a stop codon in frame with YFP in the pVYFPC-2 plasmid^1^. First, the RNaseL gene was amplified by PCR from the NA1000 genome using the following DNA primers:

1) F_01143	cgaggaaacgcatatgaaacgcctgatcgctctcaccg

2) R_01143	gcgtaacgttcgaattcgctttcttcttggcggcctgg

Next, the resulting PCR product was digested with NdeI and EcoRI enzymes, gel purified, then ligated to NdeI/EcoRI cut pVYFPC-2 plasmid using T4 DNA ligase enzyme. The ligation was transformed into chemically competent E. coli cells and selected on LB-Kan plates. The resulting kanR colonies were then screened by PCR for the insert and verified by sanger sequencing (genewiz).

**RNaseP-YFP** fusion was generated by cloning the RNaseP gene (CCNA_00807) lacking a stop codon in frame with YFP in the pVYFPC-2 plasmid^1^. First, the RNaseP gene was amplified by PCR from the NA1000 genome using the following DNA primers:

1) F_00807	cgaggaaacgcatatggctgaagcgccgcacacc

2) R_00807	cgtaacgttcgaattcgcaccggaaactgtgggatcggg

Next, the resulting PCR product was digested with NdeI and EcoRI enzymes, gel purified, then ligated to NdeI/EcoRI cut pVYFPC-2 plasmid using T4 DNA ligase enzyme. The ligation was transformed into chemically competent E. coli cells and selected on LB-Kan plates. The resulting kanR colonies were then screened by PCR for the insert and verified by sanger sequencing (genewiz).

**RNaseT2-YFP** fusion was generated by cloning the RNaseT2 gene (CCNA_00030) lacking a stop codon in frame with YFP in the pVYFPC-2 plasmid^1^. First, the RNaseT2 gene was amplified by PCR from the NA1000 genome using the following DNA primers:

1) F_00030	gcgaggaaacgcatatgttggagtcgccgatgaagaccg

2) R_00030	cgtaacgttcgaattcgcctgctggcctggtgagg

Next, the resulting PCR product was digested with NdeI and EcoRI enzymes, gel purified, then ligated to NdeI/EcoRI cut pVYFPC-2 plasmid using T4 DNA ligase enzyme. The ligation was transformed into chemically competent E. coli cells and selected on LB-Kan plates. The resulting kanR colonies were then screened by PCR for the insert and verified by sanger sequencing (genewiz).

**RNaseIII-YFP** fusion was generated by cloning the RNaseIII gene (CCNA_01630) lacking a stop codon in frame with YFP in the pVYFPC-2 plasmid^1^. First, the RNaseIII gene was amplified by PCR from the NA1000 genome using the following DNA primers:

1) F_01630	cgaggaaacgcatatggatagacgggtcgccgcc

2) R_01630	cgtaacgttcgaattcgcgcccgccccttcacgc

Next, the resulting PCR product was digested with NdeI and EcoRI enzymes, gel purified, then ligated to NdeI/EcoRI cut pVYFPC-2 plasmid using T4 DNA ligase enzyme. The ligation was transformed into chemically competent E. coli cells and selected on LB-Kan plates. The resulting kanR colonies were then screened by PCR for the insert and verified by sanger sequencing (genewiz).

**RNaseL1-YFP** fusion was generated by cloning the RNaseL1 gene (CCNA_02241) lacking a stop codon in frame with YFP in the pVYFPC-2 plasmid^1^. First, the RNaseL1 gene was amplified by PCR from the NA1000 genome using the following DNA primers:

1) F_02241	gcgaggaaacgcatatggtgtgcatgtcgaccgtcg

2) R_02241	cgtaacgttcgaattcgcgatgatctccaccaccgcg

Next, the resulting PCR product was digested with NdeI and EcoRI enzymes, gel purified, then ligated to NdeI/EcoRI cut pVYFPC-2 plasmid using T4 DNA ligase enzyme. The ligation was transformed into chemically competent E. coli cells and selected on LB-Kan plates. The resulting kanR colonies were then screened by PCR for the insert and verified by sanger sequencing (genewiz).

**RNaseR-YFP** fusion was generated by cloning the RNaseR gene (CCNA_02537) lacking a stop codon in frame with YFP in the pVYFPC-2 plasmid^1^. First, the RNaseR gene was amplified by PCR from the NA1000 genome using the following DNA primers:

1) F_02537	cgaggaaacgcatatgatggccaaacttcgccccacc

2) R_02537	gtaacgttcgaattcccgccgcttcccacgccg

Next, the resulting PCR product was digested with NdeI and EcoRI enzymes, gel purified, then ligated to NdeI/EcoRI cut pVYFPC-2 plasmid using T4 DNA ligase enzyme. The ligation was transformed into chemically competent E. coli cells and selected on LB-Kan plates. The resulting kanR colonies were then screened by PCR for the insert and verified by sanger sequencing (genewiz).

**FabG-YFP** fusion was generated by cloning the FabG gene (CCNA_00545) lacking a stop codon in frame with YFP in the pVYFPC-2 plasmid^1^. First, the FabG gene was amplified by PCR from the NA1000 genome using the following DNA primers:

1) F_00545	cgaggaaacgcatatgacgagagttgcgttcgtgaccg

2) R_00545	cgttcgaattcgcggccatgtactggccacc

Next, the resulting PCR product was digested with NdeI and EcoRI enzymes, gel purified, then ligated to NdeI/EcoRI cut pVYFPC-2 plasmid using T4 DNA ligase enzyme. The ligation was transformed into chemically competent E. coli cells and selected on LB-Kan plates. The resulting kanR colonies were then screened by PCR for the insert and verified by sanger sequencing (genewiz).

**MetK-YFP** fusion was generated by cloning the MetK gene (CCNA_00048) lacking a stop codon in frame with YFP in the pVYFPC-2 plasmid^1^. First, the MetK gene was amplified by PCR from the NA1000 genome using the following DNA primers:

1) F_00048	cgaggaaacgcatttgagccgttcgtcctacatcttcacc

2) R_00048	cgtaacgttcgaattcgccgctaggcccttcaggtcg

Next, the resulting PCR product was digested with NdeI and EcoRI enzymes, gel purified, then ligated to NdeI/EcoRI cut pVYFPC-2 plasmid using T4 DNA ligase enzyme. The ligation was transformed into chemically competent E. coli cells and selected on LB-Kan plates. The resulting kanR colonies were then screened by PCR for the insert and verified by sanger sequencing (genewiz).

N**usG-YFP** fusion was generated by cloning the NusG gene (CCNA_03310) lacking a stop codon in frame with YFP in the pVYFPC-2 plasmid^1^. First, the NusG gene was amplified by PCR from the NA1000 genome using the following DNA primers:

1) F_03310	cgaggaaacgcatatgagcaccgagaccgcg

2) R_03310	cgtaacgttcgaattcgcggcgatcttttcgacctgattgtattcc

Next, the resulting PCR product was digested with NdeI and EcoRI enzymes, gel purified, then ligated to NdeI/EcoRI cut pVYFPC-2 plasmid using T4 DNA ligase enzyme. The ligation was transformed into chemically competent E. coli cells and selected on LB-Kan plates. The resulting kanR colonies were then screened by PCR for the insert and verified by sanger sequencing (genewiz).

**RHF-YFP** fusion was generated by cloning the RHF gene (CCNA_03711) lacking a stop codon in frame with YFP in the pVYFPC-2 plasmid^1^. First, the RHF gene was amplified by PCR from the NA1000 genome using the following DNA primers:

1) F_03711	tgcgaggaaacgcatatgcaagtccaagtctccggca

2) R_03711	taacgttcgaattcgcgctcgcggttgatccgt

Next, the resulting PCR product was digested with NdeI and EcoRI enzymes, gel purified, then ligated to NdeI/EcoRI cut pVYFPC-2 plasmid using T4 DNA ligase enzyme. The ligation was transformed into chemically competent E. coli cells and selected on LB-Kan plates. The resulting kanR colonies were then screened by PCR for the insert and verified by sanger sequencing (genewiz).

**TyrRS-YFP** fusion was generated by cloning the TyrRS gene (CCNA_01946) lacking a stop codon in frame with YFP in the pVYFPC-2 plasmid^1^. First, the TyrRS gene was amplified by PCR from the NA1000 genome using the following DNA primers:

1) F_01946	gcgaggaaacgcatatgaacccctcacacaccgacc

2) R_01946	cgtaacgttcgaattcgcaacaggcttcaccaacacg

Next, the resulting PCR product was digested with NdeI and EcoRI enzymes, gel purified, then ligated to NdeI/EcoRI cut pVYFPC-2 plasmid using T4 DNA ligase enzyme. The ligation was transformed into chemically competent E. coli cells and selected on LB-Kan plates. The resulting kanR colonies were then screened by PCR for the insert and verified by sanger sequencing (genewiz).

**HFQ-YFP** fusion was generated by GenScript as a gblock fragment. The gbock fragment was cloned into pVYFPC-4 vector by GenScript.

**gblock fragment.**

cgccgaaccacgatgcgaggaaacgcatatgtccgccgaaaagaagcaaaatcttcaggacaccttcctgaacagcgttcgcaagtcgaagacgccgctcaccatcttcctcgtcaatggcgtcaagctgcagggcgtggtcagctggttcgacaatttctgcgtcctgctgcgtcgcgacggccagtcgcaactggtctacaagcacgccatttccaccatcatgccggctcaacccgttcagctctatgagccgagcgccgacgccgacgacgcgaattcgaacgttacgcgtcaccggtcggcc

**pBX-RhlE-YFP-2** plasmid was generated by subcloning the RhlE-YFP fragment from pVRhlE-YFPC-2 into pBXMCS-2 plasmid^1^. First, pVRhlE-YFPC-2 was cut with Nde1 and Nhe1 enzymes and pBXMCS-2 was cut with Nde1 and Xba1 enzymes. Next, the digested fragments were gel purified, then ligated using T4 DNA ligase enzyme. The ligation was transformed into chemically competent E. coli cells and selected on LB-Kan plates. The resulting kanR colonies were then screened by PCR for the insert and verified by sanger sequencing (genewiz).

**pBX-MetK-YFP-2** plasmid was generated by subcloning the MetK-YFP fragment from pVMetK-YFPC-2 into pBXMCS-2^1^ plasmid. First, pVMetK-YFPC-2 was cut with Nde1 and Nhe1 enzymes and pBXMCS-2 was cut with Nde1 and Xba1 enzymes. Next, the digested fragments were gel purified, then ligated using T4 DNA ligase enzyme. The ligation was transformed into chemically competent E. coli cells and selected on LB-Kan plates. The resulting kanR colonies were then screened by PCR for the insert and verified by sanger sequencing (genewiz).

**pBX-S1-2** plasmid was generated by cloning the Ribosomal Protein S1 gene (CCNA_03702) in the pBXMCS-2 plasmid^1^. First, the Ribosomal Protein S1 gene was amplified by PCR from the NA1000 genome using the following DNA primers:

1) IW002_22 RPS1_F_Nde1	ggagacgaccatatgatggctgacgatatgagcttca

2) IW003_22 RPS1_R_Xba1	ggccgctctagattagtccttggaggcccgctcgcg

Next, the resulting PCR product was digested with NdeI and Xba1 enzymes, gel purified, then ligated to NdeI/Xba1 cut pBXMCS-2 plasmid using T4 DNA ligase enzyme. The ligation was transformed into chemically competent E. coli cells and selected on LB-Kan plates. The resulting kanR colonies were then screened by PCR for the insert and verified by sanger sequencing (genewiz).

**pBX-MetK-2** plasmid was generated from pBX-MetK-YFP-2 plasmid by the inverse PCR using the following primers.

1) IW007_22 metK_F	taagctagagcggccgccaccg

2) IW006_22 metK_R	cgctaggcccttcaggtcgcccaccagat

First, the fragment was PCR amplified using the T4 PNK kinased oligos and using pBX-MetK-YFP-2 plasmid as a template. The PCR product was then DPNI treated to cut the template DNA using DPNI enzyme (Thermoscientific 10 U/μl). The DPNI treated sample was gel purified and self-ligated. The ligation was transformed into E. coli cells and selected on LB-kan plates. The resulting kan^R^ colonies were then screened by PCR for the insert and verified by sanger sequencing (genewiz).

***Strain construction***

**JS370	NA1000 vanA::papI-YFP Kan^R^**

The purified pVPAPI-YFPC-2 plasmid was transformed into NA1000 cells via electroporation and plated on PYE-Kan plates. The resulting colonies were grown in the presence and absence of vanillate and verified by fluorescence microscopy and further verified by integration PCR.

**JS371	NA1000 vanA::rhlE-YFP Kan^R^**

The purified pVRhlE-YFPC-2 plasmid was transformed into NA1000 cells via electroporation and plated on PYE-Kan plates. The resulting colonies were grown in the presence and absence of vanillate and verified by fluorescence microscopy and further verified by integration PCR.

**JS372	NA1000 vanA::rho-YFP Kan^R^**

The purified pVRhO-YFPC-2 plasmid was transformed into NA1000 cells via electroporation and plated on PYE-Kan plates. The resulting colonies were grown in the presence and absence of vanillate and verified by fluorescence microscopy and further verified by integration PCR.

**JS375	NA1000 vanA::rppH-YFP Kan^R^**

The purified pVRppH-YFPC-2 plasmid was transformed into NA1000 cells via electroporation and plated on PYE-Kan plates. The resulting colonies were grown in the presence and absence of vanillate and verified by fluorescence microscopy and further verified by integration PCR.

**JS382	NA 10000 vanA::nudC-YFP Kan^R^**

The purified pVNudC-YFPC-2 plasmid was transformed into NA1000 cells via electroporation and plated on PYE-Kan plates. The resulting colonies were grown in the presence and absence of vanillate and verified by fluorescence microscopy and further verified by integration PCR.

**JS384	NA 10000 vanA::pnp-YFP Kan^R^**

The purified pVPNP-YFPC-2 plasmid was transformed into NA1000 cells via electroporation and plated on PYE-Kan plates. The resulting colonies were grown in the presence and absence of vanillate and verified by fluorescence microscopy and further verified by integration PCR.

**JS399	NA 10000 vanA::rhlB-YFP Kan^R^**

The purified pVRhlB-YFPC-2 plasmid was transformed into NA1000 cells via electroporation and plated on PYE-Kan plates. The resulting colonies were grown in the presence and absence of vanillate and verified by fluorescence microscopy and further verified by integration PCR.

**JS410	NA 10000 vanA::acnA-YFP Kan^R^**

The purified pVAconA-YFPC-2 plasmid was transformed into NA1000 cells via electroporation and plated on PYE-Kan plates. The resulting colonies were grown in the presence and absence of vanillate and verified by fluorescence microscopy and further verified by integration PCR.

**JS431	NA 10000 vanA::rnaseJ-YFP Kan^R^**

The purified pVRNaseJ-YFPC-2 plasmid was transformed into NA1000 cells via electroporation and plated on PYE-Kan plates. The resulting colonies were grown in the presence and absence of vanillate and verified by fluorescence microscopy and further verified by integration PCR.

**JS455	NA 10000 vanA::rnaseD1-YFP Kan^R^**

The purified pVRNaseD1-YFPC-2 plasmid was transformed into NA1000 cells via electroporation and plated on PYE-Kan plates. The resulting colonies were grown in the presence and absence of vanillate and verified by fluorescence microscopy and further verified by integration PCR.

**JS481	NA 10000 vanA::rnaseHI-YFP Kan^R^**

The purified pVRNaseHI-YFPC-2 plasmid was transformed into NA1000 cells via electroporation and plated on PYE-Kan plates. The resulting colonies were grown in the presence and absence of vanillate and verified by fluorescence microscopy and further verified by integration PCR.

**JS482	NA 10000 vanA::rnaseHII-YFP Kan^R^**

The purified pVRNaseHII-YFPC-2 plasmid was transformed into NA1000 cells via electroporation and plated on PYE-Kan plates. The resulting colonies were grown in the presence and absence of vanillate and verified by fluorescence microscopy and further verified by integration PCR.

**JS483	NA 10000 vanA::smpb-YFP Kan^R^**

The purified pVSmpB-YFPC-2 plasmid was transformed into NA1000 cells via electroporation and plated on PYE-Kan plates. The resulting colonies were grown in the presence and absence of vanillate and verified by fluorescence microscopy and further verified by integration PCR.

**JS506	NA1000 vanA::rnaseL-YFP Kan^R^**

The purified pVRNaseL-YFPC-2 plasmid was transformed into NA1000 cells via electroporation and plated on PYE-Kan plates. The resulting colonies were grown in the presence and absence of vanillate and verified by fluorescence microscopy and further verified by integration PCR.

**JS508	NA1000 vanA:: rnaseP-YFP Kan^R^**

The purified pVRNaseP-YFPC-2 plasmid was transformed into NA1000 cells via electroporation and plated on PYE-Kan plates. The resulting colonies were grown in the presence and absence of vanillate and verified by fluorescence microscopy and further verified by integration PCR.

**JS510	NA1000 vanA::rnaseT2-YFP Kan^R^**

The purified pVRNaseT2-YFPC-2 plasmid was transformed into NA1000 cells via electroporation and plated on PYE-Kan plates. The resulting colonies were grown in the presence and absence of vanillate and verified by fluorescence microscopy and further verified by integration PCR.

**JS517	NA1000 vanA:: rnaseIII-YFP Kan^R^**

The purified pVRNaseIII-YFPC-2 plasmid was transformed into NA1000 cells via electroporation and plated on PYE-Kan plates. The resulting colonies were grown in the presence and absence of vanillate and verified by fluorescence microscopy and further verified by integration PCR.

**JS543	NA1000 vanA:: rnaseL1-YFP Kan^R^**

The purified pVRNaseL1-YFPC-2 plasmid was transformed into NA1000 cells via electroporation and plated on PYE-Kan plates. The resulting colonies were grown in the presence and absence of vanillate and verified by fluorescence microscopy and further verified by integration PCR.

**JS560	NA1000 vanA::rnaseR-YFP Gent^R^**

The purified pVRNaseR-YFPC-4 plasmid was transformed into NA1000 cells via electroporation and plated on PYE-Gent plates. The resulting colonies were grown in the presence and absence of vanillate and verified by fluorescence microscopy and further verified by integration PCR.

**JS678	NA1000 vanA::fabG-YFP Kan^R^**

The purified pVFabG-YFPC-2 plasmid was transformed into NA1000 cells via electroporation and plated on PYE-Kan plates. The resulting colonies were grown in the presence and absence of vanillate and verified by fluorescence microscopy and further verified by integration PCR.

**JS680	NA1000 vanA::metK-YFP Kan^R^**

The purified pVMetK-YFPC-2 plasmid was transformed into NA1000 cells via electroporation and plated on PYE-Kan plates. The resulting colonies were grown in the presence and absence of vanillate and verified by fluorescence microscopy and further verified by integration PCR.

**JS688	NA1000 vanA::nusG-YFPC Kan^R^**

The purified pVNusG-YFPC-2 plasmid was transformed into NA1000 cells via electroporation and plated on PYE-Kan plates. The resulting colonies were grown in the presence and absence of vanillate and verified by fluorescence microscopy and further verified by integration PCR.

**JS690	NA1000 vanA::rhf-YFP Kan^R^**

The purified pV-RHF-YFPC-2 plasmid was transformed into NA1000 cells via electroporation and plated on PYE-Kan plates. The resulting colonies were grown in the presence and absence of vanillate and verified by fluorescence microscopy and further verified by integration PCR.

**JS692	NA1000 vanA::tyrRS-YFP Kan^R^**

The purified pVTyrRS-YFPC-2 plasmid was then transformed into NA1000 cells via electroporation and plated on PYE-Kan plates. The resulting colonies were grown in the presence and absence of vanillate and verified to be expressing the YFP fusion by fluorescence microscopy and further verified by integration PCR.

**JS697	NA1000 vanA::hfq-YFP Gent^R^**

The purified pVHFQ-YFPC-4 plasmid was transformed into NA1000 cells via electroporation and plated on PYE-Gent plates. The resulting colonies were grown in the presence and absence of vanillate and verified by fluorescence microscopy and further verified by integration PCR.

**JS699 NA1000 vanA::hfq-YFPC rne::pXrnessrAC Gent^R^ Kan^R^**

This strain was generated by transducing JS697 cells with RNase E depletion phage lysate from JS8^2^ strain. The transduced cells were plated in PYE-Gent-Kan-Xylose plates and the resultant colonies were grown in PYE-Gent-Kan media with and without xylose to verify the RNase E depletion phenotype.

**JS694 NA1000 rne::pXrnessrAC Gent^R^**

The plasmid pXMCS-4^1^ containing last 500bp of RNase E gene was transformed into NA1000 cells via electroporation and the colonies were selected on PYE-Gent-Xylose plates. The integration of RNase E gene under the xylose inducible promoter was confirmed by growing the cells in the presence and absence of xylose

**JS701 NA1000 vanA::rhlE-YFPC rne::pXrnessrAC Gent^R^ Kan^R^**

This strain was generated by transducing JS371 cells with RNase E depletion phage lysate from JS694 strain. The transduced cells were plated in PYE-Gent-Kan-Xylose plates and the resultant colonies were grown in PYE-Gent-Kan media with and without xylose to verify the RNase E depletion phenotype.

**JS702 NA1000 vanA::rho-YFPC rne::pXrnessrAC Gent^R^ Kan^R^**

This strain was generated by transducing JS372 cells with RNase E depletion phage lysate from JS694 strain. The transduced cells were plated in PYE-Gent-Kan-Xylose plates and the resultant colonies were grown in PYE-Gent-Kan media with and without xylose to verify the RNase E depletion phenotype.

**JS706 NA1000 vanA::metK-YFPC rne::pXrnessrAC Gent^R^ Kan^R^**

This strain was generated by transducing JS680 cells with RNase E depletion phage lysate from JS694 strain. The transduced cells were plated in PYE-Gent-Kan-Xylose plates and the resultant colonies were grown in PYE-Gent-Kan media with and without xylose to verify the RNase E depletion phenotype.

**JS709 NA1000 vanA::rnaseD1(CTD)-YFPC rne::pXrnessrAC Gent^R^ Kan^R^**

This strain was generated by transducing JS397 cells with RNase E depletion phage lysate from JS694 strain. The transduced cells were plated in PYE-Gent-Kan-Xylose plates and the resultant colonies were grown in PYE-Gent-Kan media with and without xylose to verify the RNase E depletion phenotype.

**JS451 NA1000 acnA::acnA-Chy pBXMCS Spec^R^ Strp^R^ Kan^R^**

The strain was generated by electroporation of JS134^2^ with pBXMCS-2 plasmid^1^. The cells were plated on PYE-Spec-Strp-Kan plates. The resulting colonies were grown in the presence and absence of xylose and verified by fluorescence microscopy.

**JS452 NA1000 acnA::acnA-Chy rne::pBXrne-YFPC Spec^R^ Strp^R^ Kan^R^**

The strain was generated by electroporation of JS134^2^ with pBXRNE-YFP-2 plasmid^3^. The cells were plated on PYE-Spec-Strp-Kan plates. The resulting colonies were grown in the presence and absence of xylose and verified by fluorescence microscopy.

**JS716	NA1000 acnA::acnA-Chy rhlE::pBXrhlE-YFPC Spec^R^ Strp^R^ Kan^R^**

The strain was generated by electroporation of JS134^2^ with pBX-RhlE-YFP-2 plasmid. The cells were plated on PYE-Spec-Strp-Kan plates. The resulting colonies were grown in the presence and absence of xylose and verified by fluorescence microscopy.

**JS721	NA1000 acnA::acnA-Chy RPS1::pBXRPS1 Spec^R^ Strp^R^ Kan^R^**

The strain was generated by electroporation of JS134^2^ with pBX-S1-2 plasmid. The cells were plated on PYE-Spec-Strp-Kan plates. The resulting colonies were grown in the presence and absence of xylose and verified by fluorescence microscopy.

**JS742**	**NA1000 acnA::acnA-Chy metK::pBXmetK Spec^R^ Strp^R^ Kan^R^**

The strain was generated by electroporation of JS134^2^ with pBX-MetK-2 plasmid. The cells were plated on PYE-Spec-Strp-Kan plates. The resulting colonies were grown in the presence and absence of xylose and verified by fluorescence microscopy.

**JS563	NA1000 vanA::rhlE-YFP rne::rne-CHYC Kan^R^ Gent^R^**

This strain was generated by transducing JS371 cells with pRNE-chy phage lysate from JS403 strain. The transduced cells were plated in PYE-Gent-Kan plates. The resulting colonies were grown in the presence and absence of vanillate and verified by fluorescence microscopy and further verified by integration PCR.

**JS564	NA1000 vanA::rho-YFP rne::rne-CHYC Kan^R^ Gent^R^**

This strain was generated by transducing JS372 cells with pRNE-chy phage lysate from JS403 strain. The transduced cells were plated in PYE-Gent-Kan plates. The resulting colonies were grown in the presence and absence of vanillate and verified by fluorescence microscopy and further verified by integration PCR.

**JS570	NA1000 vanA::rnaseHI-YFP rne::rne-CHYC Kan^R^ Gent^R^**

This strain was generated by transducing JS481 cells with pRNE-chy phage lysate from JS403 strain. The transduced cells were plated in PYE-Gent-Kan plates. The resulting colonies were grown in the presence and absence of vanillate and verified by fluorescence microscopy and further verified by integration PCR.

**JS758	NA1000 vanA::rnaseD1-YFP rne::rne-CHYC Kan^R^ Gent^R^**

This strain was generated by transducing JS455 cells with pRNE-chy phage lysate from JS403 strain. The transduced cells were plated in PYE-Gent-Kan plates. The resulting colonies were grown in the presence and absence of vanillate and verified by fluorescence microscopy and further verified by integration PCR.

**JS761	NA1000 vanA::metK-YFP rne::rne-CHYC Kan^R^ Gent^R^**

This strain was generated by transducing JS680 cells with **pRNE-chy-4 phage lysate**. The transduced cells were plated in PYE-Gent-Kan plates. The resulting colonies were grown in the presence and absence of vanillate and verified by fluorescence microscopy and further verified by integration PCR.

**JS735	NA1000 rne::rne-msfGFP Gent^R^**

This strain was generated by transducing NA1000 cells with RNase E-msfGFP phage lysate from JS87^2^ strain. The transduced cells were plated in PYE-Gent plates. The resultant colonies were grown in PYE-Gent media and verified to be expressing the RNase E-msfGFP fusion by fluorescence microscopy.

**JS744	NA1000 rne::rne(∆hfq)-msfGFP Gent^R^ Spec^R^ Strp^R^**

This strain was generated by transducing ∆*hfq* (Shapiro Lab, Stanford University School of Medicine) cells with RNase E-msfGFP phage lysate from JS87^2^ strain. The transduced cells were plated in PYE-Gent-Spec-Strp plates. The resultant colonies were grown in PYE-Gent-Spec-Strp and verified to be expressing the RNase E-msfGFP fusion by fluorescence microscopy.

**JS671	NA1000 rne::rne(∆rhlE)-msfGFP Gent^R^ Kan^R^**

This strain was generated by transducing RhlE disruption mutant (Marques Lab, University of Sao Paulo) cells with RNase E-msfGFP phage lysate from JS87^2^ strain. The transduced cells were plated in PYE-Gent-Kan plates. The resultant colonies were grown in PYE-Gent-Kan media verified to be expressing the RNase E-msfGFP fusion by fluorescence microscopy.

**JS672	NA1000 rne::rne(∆rhlB)-msfGFP Gent^R^**

This strain was generated by transducing ∆*rhlB* (Marques Lab, University of Sao Paulo) cells with RNase E-msfGFP phage lysate from JS87^2^ strain. The transduced cells were plated in PYE-Gent plates. The resultant colonies were grown in PYE-Gent media and verified to be expressing the RNase E-msfGFP fusion by fluorescence microscopy.

**JS128	NA1000 *popZ-Chy rne::pXrnessrAC* Kan^R^**

This strain was generated by transducing cells harboring popZ-mCherry (native gene fusion) with RNase E depletion phage lysate from JS82 strain. The transduced cells were plated in PYE-Kan-Xylose plates and the resultant colonies were grown in PYE-Kan media with and without xylose to verify the RNase E depletion phenotype.

**Strained used in Bacterial two hybrid assay**

**VNS003**

The strain was created by transforming pVN054 and pVN057 plasmids into BTH101 cells and plated on LB-Agar plates containing Kanamycin, Ampicillin, IPTG and X-gal. The resultant colonies were selected based on blue or white color of the colonies.

**VNS004**

The strain was created by transforming pVN054 and pVN055 plasmids into BTH101 cells and plated on LB-Agar plates containing Kanamycin, Ampicillin, IPTG and X-gal. The resultant colonies were selected based on blue or white color of the colonies.

**VNS005**

The strain was created by transforming pVN054 and pVN058 plasmids into BTH101 cells and plated on LB-Agar plates containing Kanamycin, Ampicillin, IPTG and X-gal. The resultant colonies were selected based on blue or white color of the colonies.

**VNS006**

The strain was created by transforming pKT25-Zip and pUT18C-Zip plasmids into BTH101 cells and plated on LB-Agar plates containing Kanamycin, Ampicillin, IPTG and X-gal. The resultant colonies were selected based on blue or white color of the colonies.

**VNS007**

The strain was created by transforming pKNT25 and pUT18 plasmids into BTH101 cells and plated on LB-Agar plates containing Kanamycin, Ampicillin, IPTG and X-gal. The resultant colonies were selected based on blue or white color of the colonies.

**VNS008**

The strain was created by transforming pVN054 and pUT18 plasmids into BTH101 cells and plated on LB-Agar plates containing Kanamycin, Ampicillin, IPTG and X-gal. The resultant colonies were selected based on blue or white color of the colonies.

**VNS009**

The strain was created by transforming pKNT25 and pVN055 plasmids into BTH101 cells and plated on LB-Agar plates containing Kanamycin, Ampicillin, IPTG and X-gal. The resultant colonies were selected based on blue or white color of the colonies.

**VNS010**

The strain was created by transforming pKNT25 and pVN058 plasmids into BTH101 cells and plated on LB-Agar plates containing Kanamycin, Ampicillin, IPTG and X-gal. The resultant colonies were selected based on blue or white color of the colonies.

***List of Tables***

**Table S1.** Log2 enrichment (JS299/JS221) values of all proteins (related to Figure 1).

**Table S2.** Disorderness, Pscore, Catgranule and DeePhase scores of all proteins used in *in vitro* assays (related to Figure 3).

**Table S3.** Strains used in this study.

**Table S4.** Plasmids used in this study.

**Table S5.** DNA oligos used in this study
