## Supplemental Figures for "The BR-body proteome contains a complex network of protein-protein and protein-RNA interactions"

### Figure S1

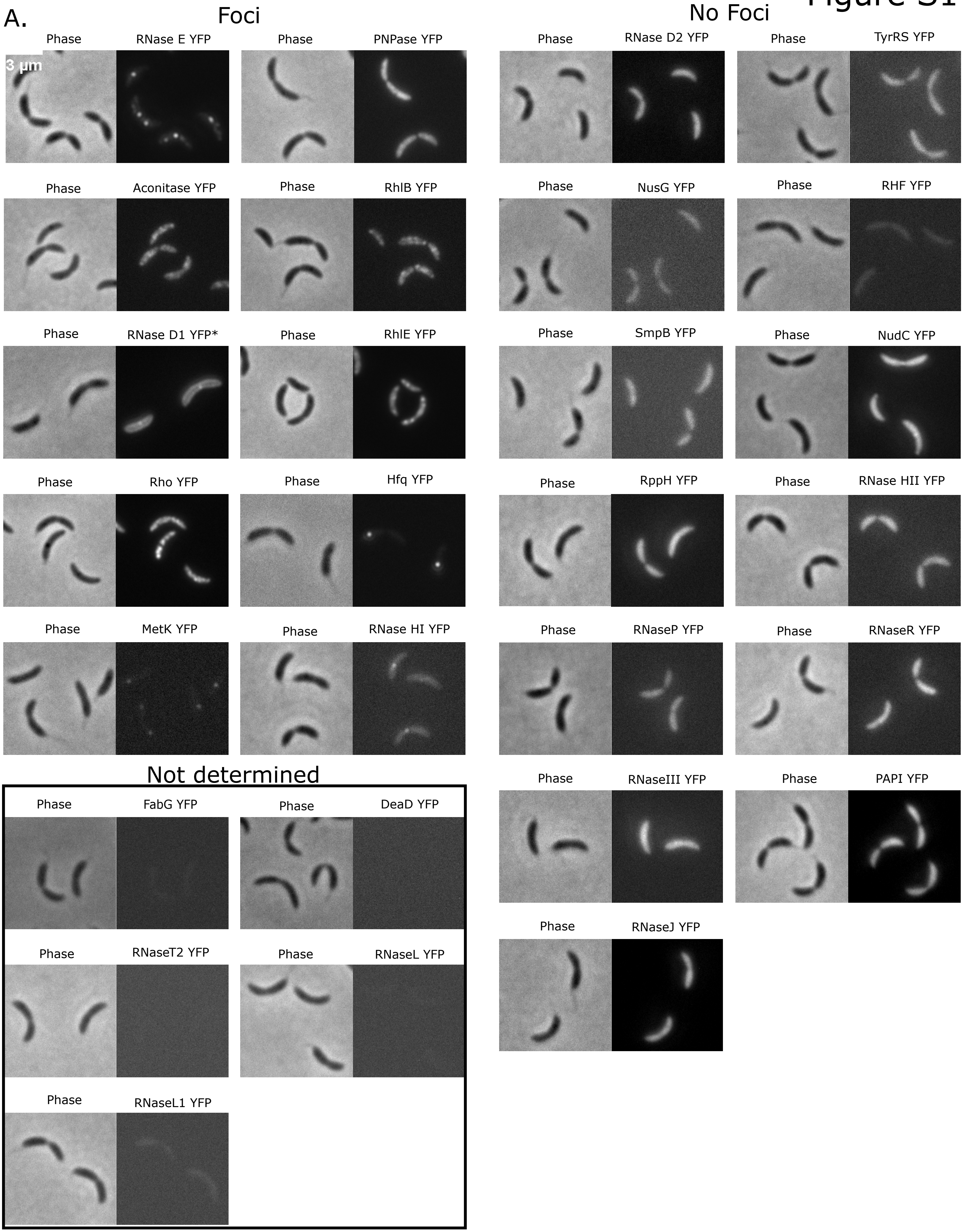
A.
Foci
No Foci
Phase
RNase E YFP
3 µm
PNPase YFP
Aconitase YFP
RhlB YFP
RNase D1 YFP*
RhlE YFP
Rho YFP
Hfq YFP
MetK YFP
RNase HI YFP
RNase D2 YFP
TyrRS YFP
NusG YFP
RHF YFP
SmpB YFP
NudC YFP
RppH YFP
RNase HII YFP
RNaseP YFP
RNaseR YFP
RNaseIII YFP
PAPI YFP
RNaseJ YFP
Not determined
FabG YFP
DeaD YFP
RNaseT2 YFP
RNaseL YFP
RNaseL1 YFP

### Figure S2

A.

Xylose Dependent RNase E Depletion strain

+Xyl

-Xyl

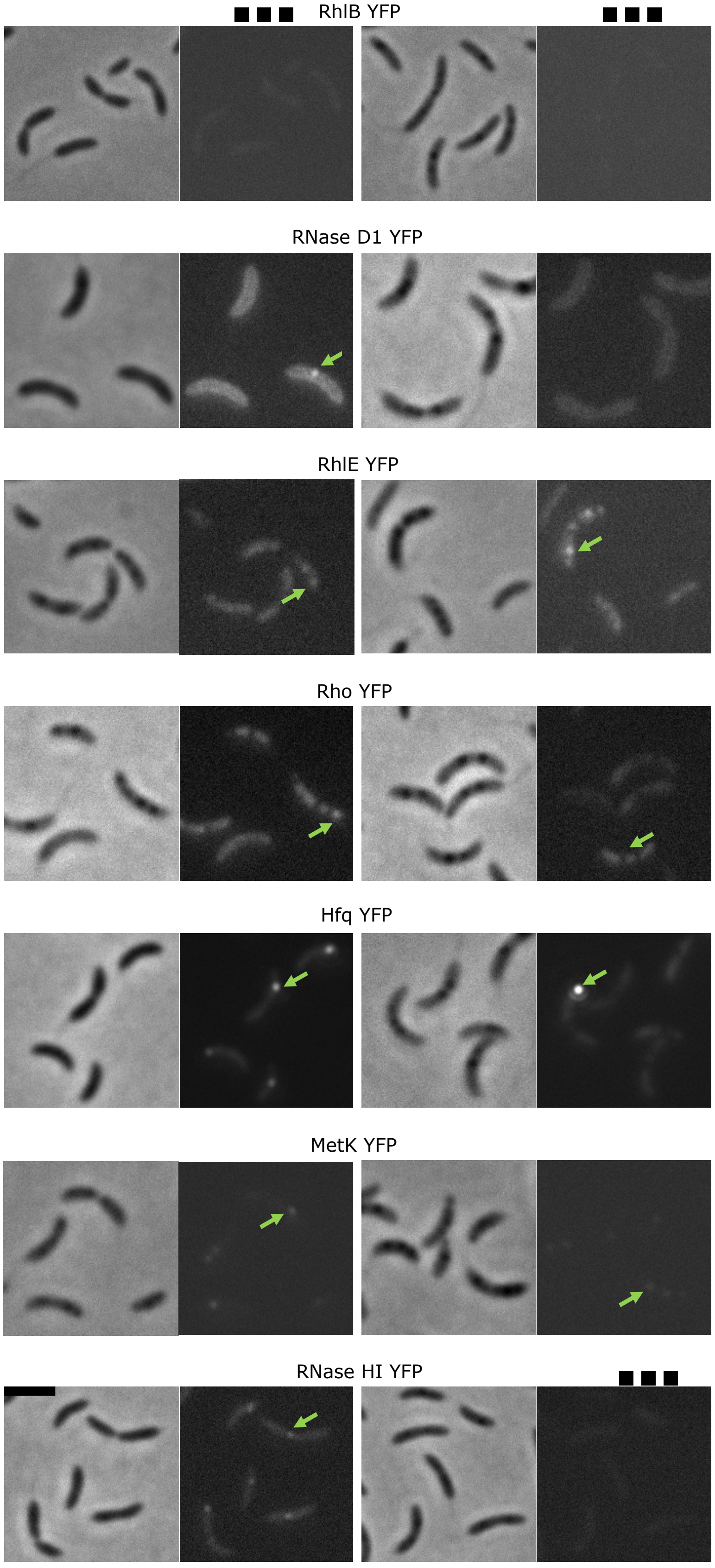

Figure S3

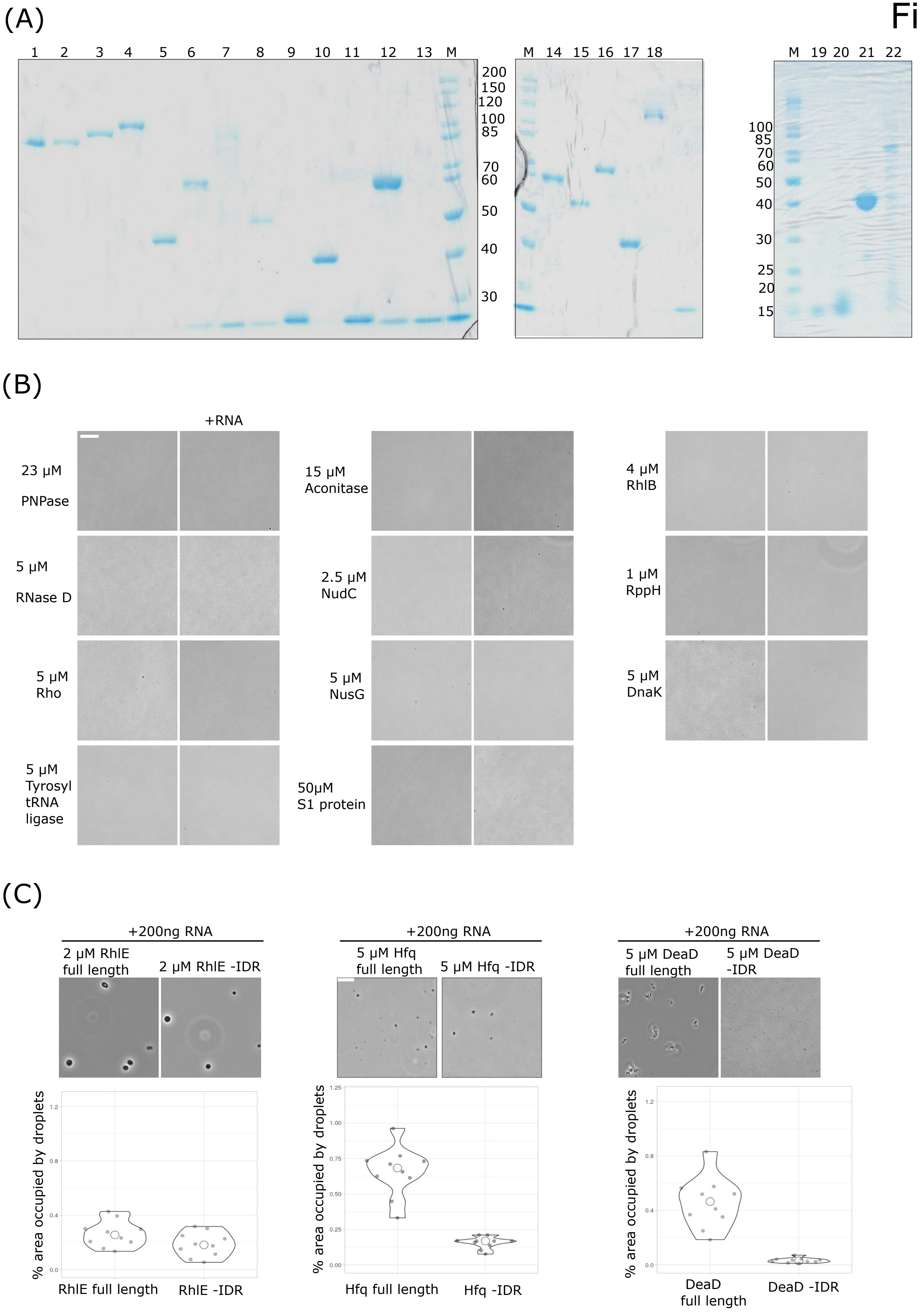

(A)
1 2 3 4 5 6 7 8 9 10 11 12 13 M
200
150
120
100
85
70
60
50
40
30
M 14 15 16 17 18
M 19 20 21 22
100
85
70
60
50
40
30
25
20
15
(B)
+RNA
23 µM PNPase
15 µM Aconitase
4 µM RhlB
5 µM RNase D
2.5 µM NudC
1 µM RppH
5 µM Rho
5 µM NusG
5 µM DnaK
5 µM Tyrosyl tRNA ligase
50µM S1 protein
(C)
+200ng RNA
2 µM RhlE full length
2 µM RhlE -IDR
5 µM Hfq full length
5 µM Hfq -IDR
5 µM DeaD full length
5 µM DeaD -IDR
% area occupied by droplets
RhlE full length
RhlE -IDR
Hfq full length
Hfq -IDR
DeaD full length
DeaD -IDR

### Figure S4

(A)

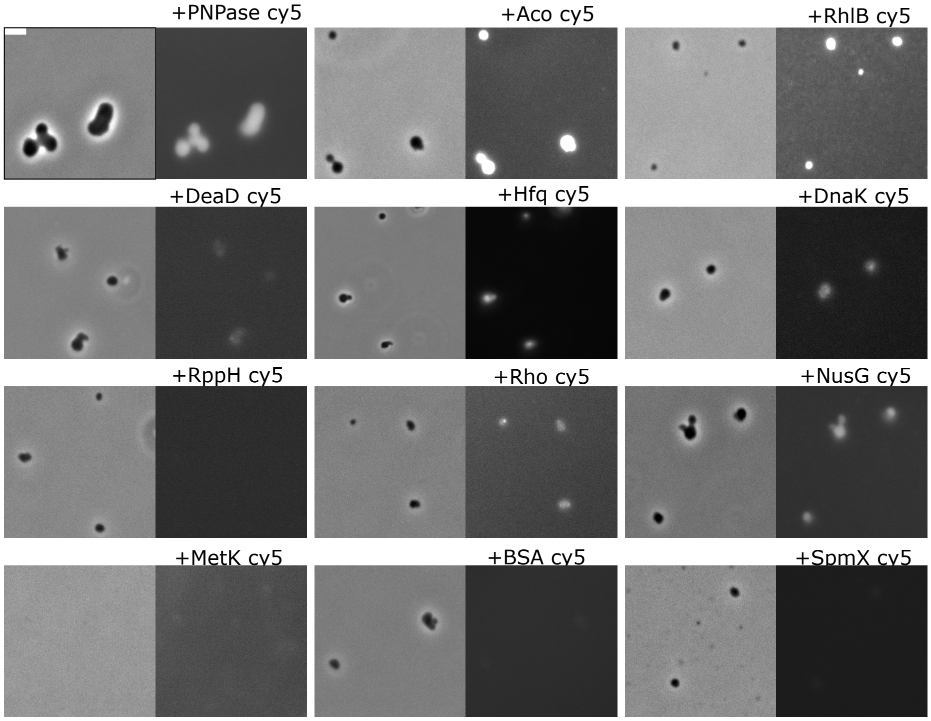

+PNPase cy5
+Aco cy5
+RhlB cy5
+RND1 cy5
+RhlE cy5
+DeaD cy5
+Hfq cy5
+DnaK cy5
+FabG cy5
+NudC cy5
+RppH cy5
+Rho cy5
+NusG cy5
+TyrRS cy5
+S1 cy5
+MetK cy5
+BSA cy5
+SpmX cy5


(B)

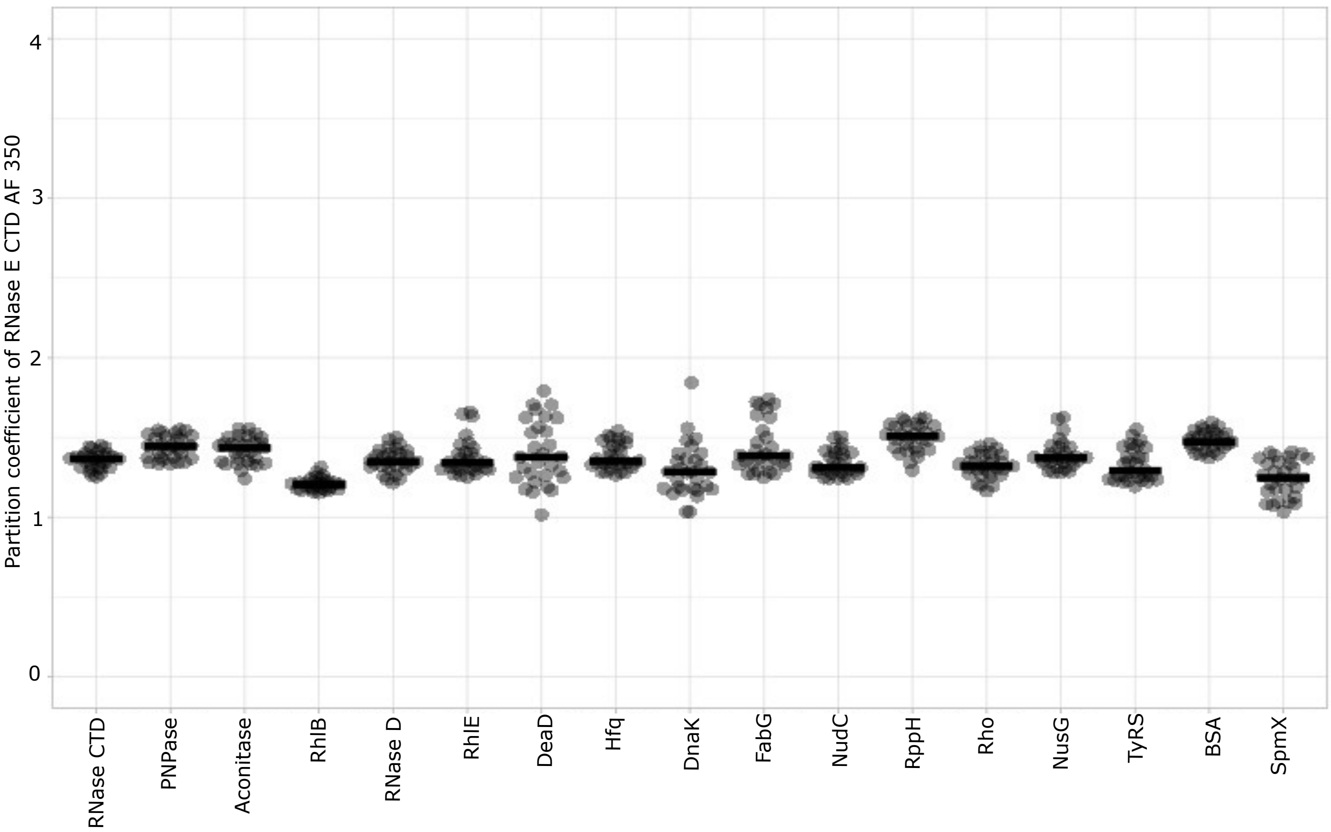

Partition coefficient of RNase E CTD AF 350
0
1
2
3
4
RNase CTD
PNPase
Aconitase
RhlB
RNase D
RhlE
DeaD
Hfq
DnaK
FabG
NudC
RppH
Rho
NusG
TyRS
BSA
SpmX

### Figure S5

A.

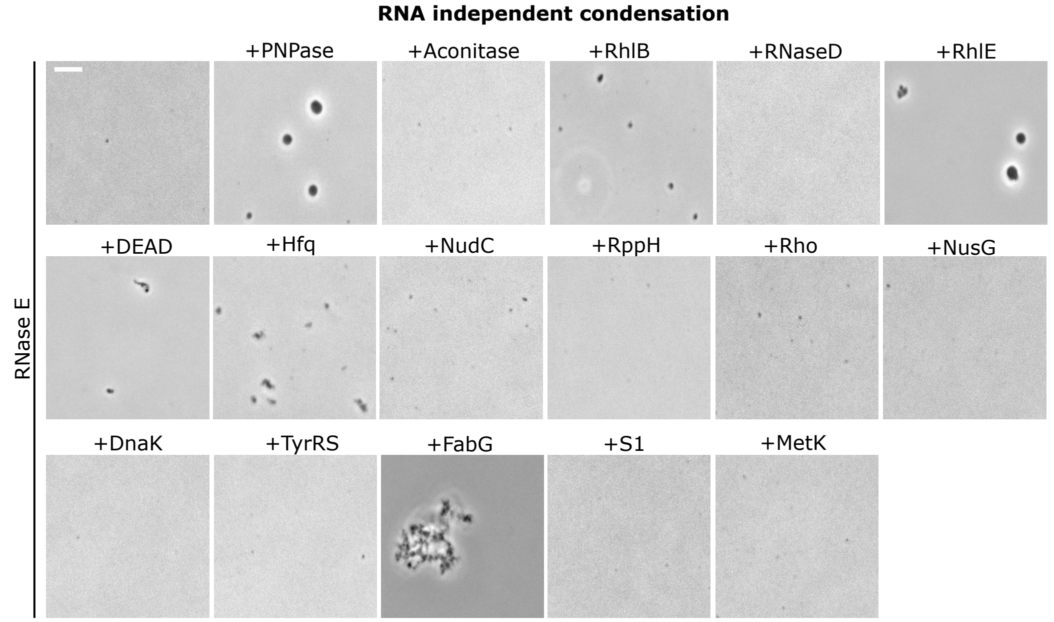


B.

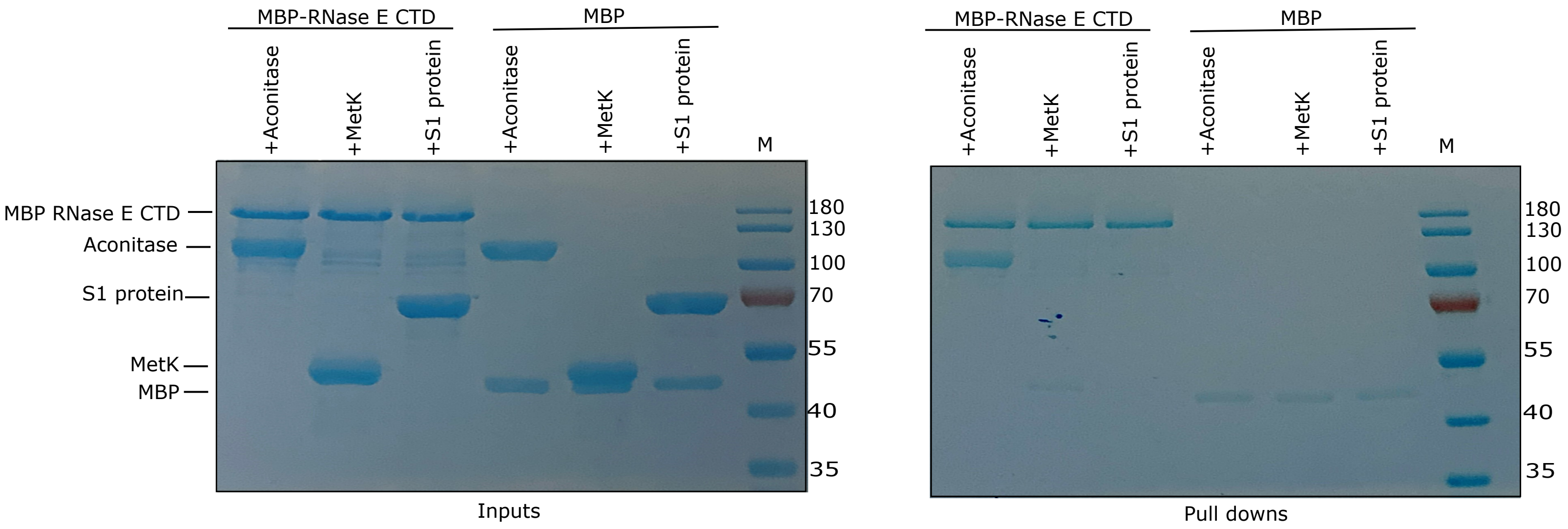


C.

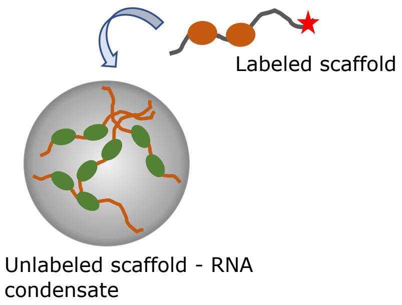


| -RNA Scaffold | Labeled Scaffold | | | | |
|---|---|---|---|---|---|
| | RNase E | RhlE | DeaD | Hfq | FabG |
| RNase E | X | 12 | 1.4 | 7.5 | 1.9 |
| RhlE | 12 | X | 9.8 | 2.3 | 2.2 |
| DeaD | 2.6 | 15 | X | 3.0 | 3.9 |
| Hfq | 6.9 | 3.7 | ND | X | 2.2 |
| FabG | ND | ND | ND | ND | X |

Figure S6

A.

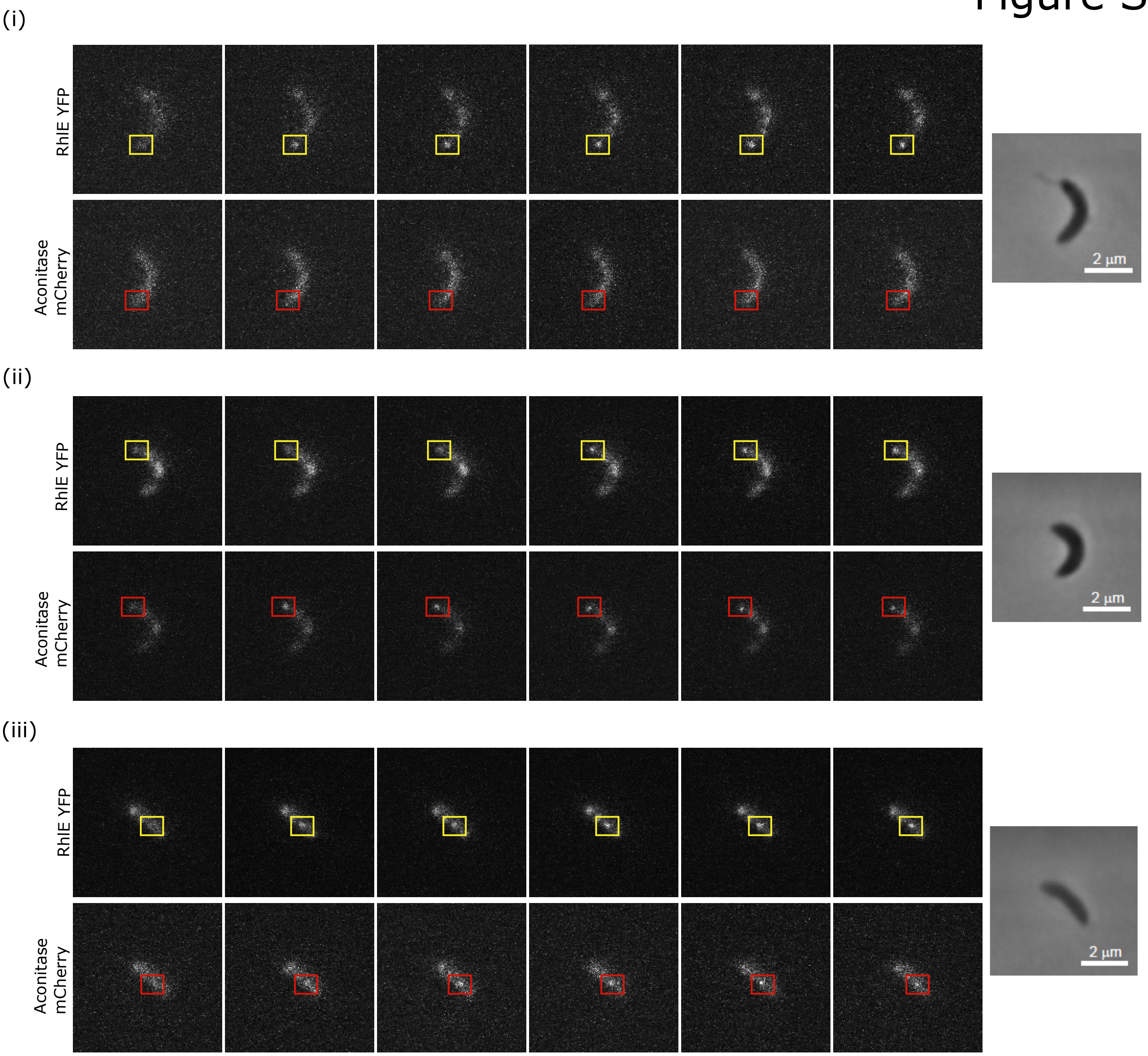

(i)
RhlE YFP
Aconitase mCherry
2 µm
(ii)
RhlE YFP
Aconitase mCherry
2 µm
(iii)
RhlE YFP
Aconitase mCherry
2 µm


B.

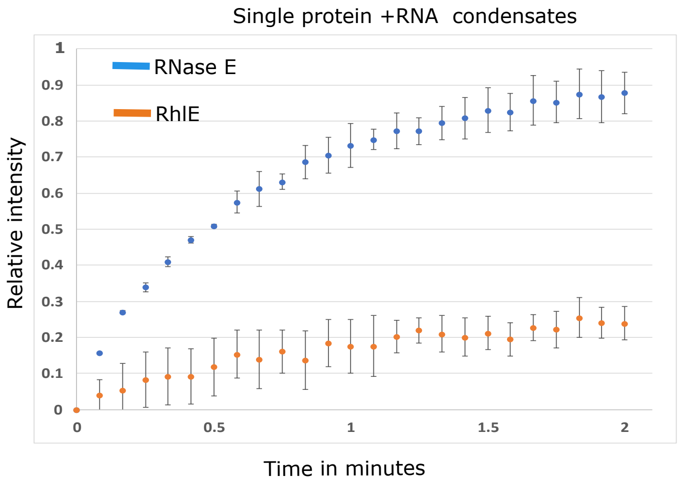

Single protein +RNA condensates
RNase E
RhlE
Relative intensity
Time in minutes
0
0.1
0.2
0.3
0.4
0.5
0.6
0.7
0.8
0.9
1
0
0.5
1
1.5
2


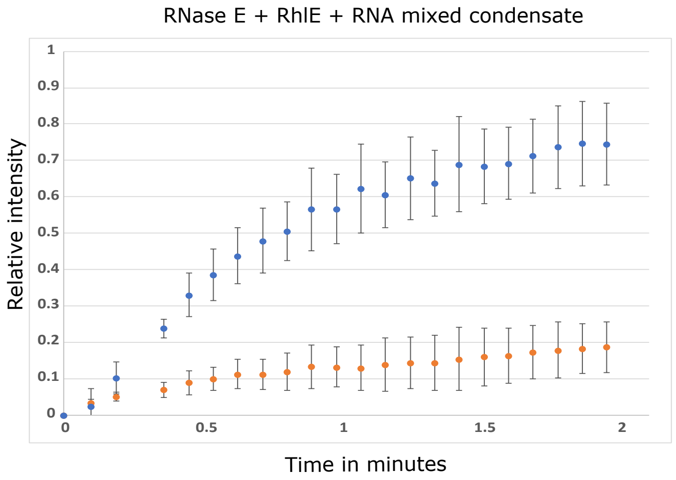

RNase E + RhlE + RNA mixed condensate
Relative intensity
Time in minutes
0
0.1
0.2
0.3
0.4
0.5
0.6
0.7
0.8
0.9
1
0
0.5
1
1.5
2

### Figure S7

6 μM RNase E - 30 mins incubation

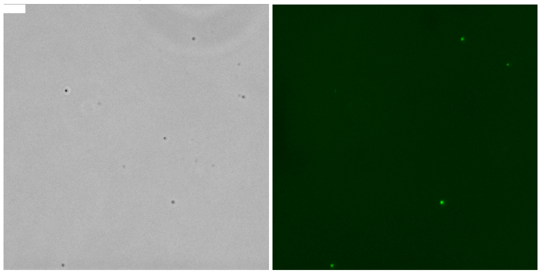

20 μM RNase E - 30 mins incubation

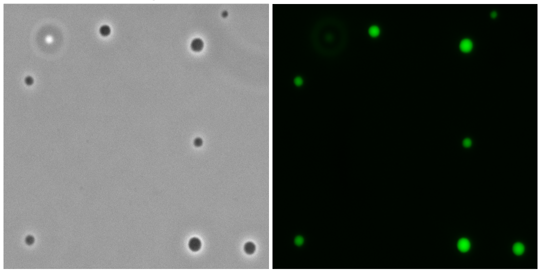
